# Marker-based CRISPR screens identify POU2F1 as a regulator of DLL3 and neuroendocrine identity in small cell lung cancer

**DOI:** 10.64898/2026.04.08.717069

**Authors:** Patrick J. Cunniff, Connor Fitzpatrick, Jack Bauer, Damianos Skopelitis, Olaf Klingbeil, Toyoki Yoshimoto, Leemor Joshua-Tor, Christopher R. Vakoc

## Abstract

Small cell lung cancers (SCLC) often exhibit a neuroendocrine lineage identity marked by high expression of Delta-like Ligand 3 (DLL3). Because DLL3 shows minimal expression in normal adult tissues, it serves as an SCLC-selective tumor antigen and is the basis for clinically efficacious targeted therapies. Understanding the mechanisms that regulate DLL3 expression is therefore critical for advancing therapeutic strategies in this disease. Here, we performed transcription factor-focused and genome-wide CRISPR screens to identify regulators of DLL3 expression in SCLC. Both approaches converged on POU2F1 as a top activator of *DLL3* in this tumor context. Despite its ubiquitous expression, we identify an SCLC-specific role for POU2F1 in activating *DLL3* and a broader set of neuroendocrine lineage genes. Epigenomic analyses reveal tandem POU2F1–ASCL1 motifs within the *DLL3* promoter that underlie the strong codependency between POU2F1 and the neuroendocrine master regulator ASCL1 for high-level DLL3 expression in SCLC. We provide evidence that tandem POU2F1–ASCL1 elements are part of a cis-regulatory code for the lung neuroendocrine cell fate. Together, these findings define a previously unrecognized transcriptional logic controlling DLL3 expression and establish POU2F1 as a context-specific regulator of neuroendocrine lineage in small cell lung cancer.

## Introduction

Small cell lung cancer (SCLC) has long been recognized as a distinct clinicopathologic entity characterized by rapid cell proliferation, early dissemination, and a robust neuroendocrine phenotype^1^. From its earliest pathological descriptions, SCLC tumors were noted to express neuroendocrine markers such as synaptophysin and chromogranin A^2^, linking this disease to neuroendocrine differentiation before the advent of molecular profiling^3^. Clinically, this lineage identity has shaped diagnostic criteria, informed therapeutic approaches, and distinguished SCLC from other lung cancer subtypes, despite shared etiologic factors such as tobacco exposure^1^. More recent genomic and transcriptomic studies have refined this view by revealing that SCLC comprises multiple molecular subtypes defined by dominant transcriptional regulators, including ASCL1, NEUROD1, and POU2F3^4^. Among these, ASCL1-positive SCLC, hereafter referred to as SCLC-A, represents the most prevalent and canonical neuroendocrine subtype^4^.

A prominent feature of the neuroendocrine state in SCLC is high expression of Delta-like ligand 3 (DLL3), an atypical member of the Notch ligand family^5^. In contrast to canonical Delta-like ligands that activate Notch signaling at the cell surface, DLL3 primarily localizes to intracellular compartments and functions as a negative regulator of Notch pathway activity during development^6,7^. DLL3 expression is largely restricted to embryonic neuroendocrine contexts and is minimally expressed in most adult tissues^8^. In SCLC, however, DLL3 is highly and selectively expressed, particularly within tumors with neuroendocrine lineage identity^9^. This restricted expression pattern has positioned DLL3 as a clinically important tumor-associated antigen and has driven the development of multiple DLL3-directed therapeutic strategies^10–16^. Beyond its translational relevance, DLL3 has emerged as a robust marker of neuroendocrine differentiation in SCLC, linking suppression of Notch signaling to maintenance of lineage identity^17^.

Despite its central role in SCLC biology and therapy, the transcriptional mechanisms that govern DLL3 expression in this disease remain incompletely defined. Prior work has largely inferred DLL3 regulation indirectly through its association with ASCL1-positive tumors and low Notch signaling, but the specific transcription factors and cis-regulatory elements responsible for DLL3 activation have not been systematically characterized^17^. This gap is notable given the growing clinical reliance on DLL3-directed therapies and the increasing recognition that SCLC exhibits substantial transcriptional plasticity^18,19^. A clearer understanding of how DLL3 expression is encoded at the level of transcriptional regulation is therefore necessary to explain its subtype specificity, predict its stability under therapeutic pressure, and to identify mechanisms by which DLL3 expression may be gained or lost during tumor evolution.

Functional genetic screening has emerged as a powerful and unbiased approach to dissect the transcriptional and lineage dependencies that underlie SCLC biology^20–23^. CRISPR-based loss-of-function screens have been particularly informative in this disease, revealing subtype-specific regulators, lineage vulnerabilities, and chromatin dependencies that are not readily inferred from gene expression or genomic data alone^23^. Using this strategy, our group previously identified POU2F3 as a master regulator of a non-neuroendocrine tuft-cell-like SCLC subtype^22^ (often referred to as SCLC-P) and uncovered a selective dependence of these tumors on specific transcriptional coactivators^21,24^, establishing a mechanistic framework for tuft cell-like identity in SCLC. Studies from other groups have similarly leveraged CRISPR screening to define core transcriptional regulators, chromatin modifiers, and signaling pathways that sustain distinct SCLC states and govern lineage plasticity^25–27^. These studies highlight the power of functional screening to identify previously unknown lineage regulators.

POU2F1, also known as OCT1, is a ubiquitously expressed POU-domain transcription factor best known for its roles in basal promoter activity, metabolism, stress responses, and signal-dependent gene regulation across diverse cell types^28–32^. Unlike lineage-restricted POU transcription factors, POU2F1 has generally been viewed as a transcription factor that supports general cellular functions and integrates environmental cues^33^. Prior studies have implicated POU2F1 in processes such as immune activation^34^, metabolic adaptation^35^, and cellular stress tolerance^36^, with context-dependent effects shaped by chromatin accessibility and cofactor availability. In a few contexts, POU2F1 has been found to regulate normal and malignant cell identity programs, often through cooperativity with other transcription factors^37–39^. However, POU2F1 has not been previously linked to neuroendocrine differentiation or to neuroendocrine lineage cancers. As a result, it remains unclear how a widely expressed transcription factor such as POU2F1 might acquire specialized functions within specific tumor lineages.

In this study, we set out to define the transcriptional mechanisms that drive DLL3 expression in SCLC. By combining two CRISPR screening approaches, we identify POU2F1 as a previously unappreciated regulator of DLL3 expression and of neuroendocrine transcriptional programs in this disease. We show that POU2F1 collaborates with ASCL1 through a defined cis-regulatory architecture at the *DLL3* promoter, providing a mechanistic explanation for how a ubiquitously expressed transcription factor can promote lineage-restricted gene expression. Extending beyond *DLL3*, our findings reveal a regulatory principle in which tandem POU2F1–ASCL1 motifs encode neuroendocrine transcriptional programs. Together, this work links lineage identity to tumor-selective antigen expression in SCLC and offers insight into how neuroendocrine cell states are maintained.

## Results

### A CRISPR screening strategy for identifying regulators of DLL3 expression in SCLC

Aberrant expression of the Notch ligand DLL3 is a defining feature of small cell lung cancer (SCLC) and underlies several therapeutic strategies, yet the mechanisms that drive DLL3 expression remain poorly understood. To identify upstream regulators of DLL3 in an unbiased manner, we developed a CRISPR screening strategy to systematically nominate genes required for DLL3 expression in SCLC. We selected the human SCLC cell line NCI-H209 for these screens based on its robust growth in culture and its high expression of both DLL3 and the neuroendocrine master regulator ASCL1 (Fig. S1A, Supplementary Table 1). We identified an antibody capable of reliably and quantitatively detecting DLL3 protein and validated it by Western blotting and flow cytometry, confirming its specificity using CRISPR-mediated inactivation of *DLL3* as a negative control (Figs. 1A–B).

**Figure 1.**
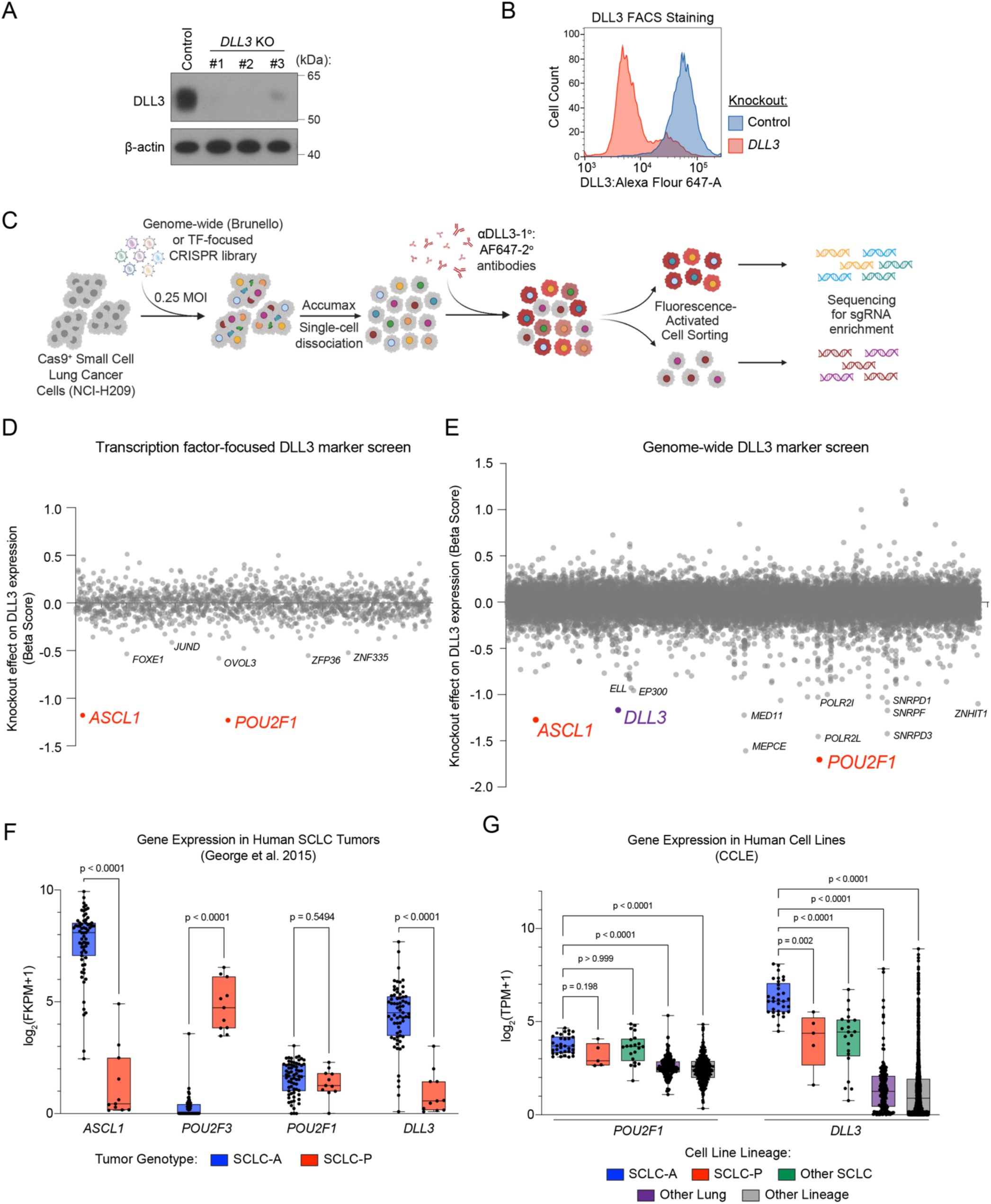
Parallel marker-based screens identify POU2F1 as a DLL3 regulator. (A) Western blot of DLL3 protein 7 days after CRISPR-Cas9 knockout (KO) of *DLL3* or *ROSA26* (control) in H209 cells. β-actin, loading control. Representative of two biological replicates. (B) Flow cytometry analysis of H209 cells, methanol-fixed and stained with anti-DLL3 antibodies 8 days following KO of *DLL3* or control (*ROSA26*). Representative of 2 biological replicates. (C) Workflow of genome-wide Serpin B5 reporter screen. Created in BioRender. Cunniff, P. (2026) https://BioRender.com/l9r3z2o. (D–E) Transcription factor-focused (D) or Genome-wide (E) DLL3 reporter screen results in H209. Genes (dots) ordered alphabetically along the x-axis. Select outlier genes are labeled. Beta scores and significance calculated using MAGeCK (maximum likelihood estimation). Negative beta scores indicate enrichment in the DLL3^low^ population. (F–G) Gene expression in (F) resected human small cell lung cancer tumors – reanalyzed from George et al. (2015) or (G) human cell lines. Each point = 1 tumor or cell line. Boxes represent the mean ± interquartile range (IQR). Error bars extend to the minimum and maximum values. Statistical significance was evaluated using two-sided Mann–Whitney tests with (F) no multiple comparison correction or (G) correction for 4 comparisons. TPM = transcripts per million.

For the CRISPR screens, Cas9-expressing NCI-H209 cells were transduced with two different lentiviral sgRNA libraries. The first library targets 1,437 DNA-binding domains (DBDs) to enable efficient coverage of the human transcription factor repertoire (8,739 sgRNAs total)^22^. The second was a genome-wide CRISPR sgRNA library targeting 19,115 human genes (Brunello)^40^. On day 8 following transduction, 50 (DBD) or 750 (Brunello) million cells were harvested, dissociated to single cells, methanol-fixed, and stained with anti-DLL3 primary and fluorophore-conjugated secondary antibodies. Cells were then sorted by flow cytometry into populations with low, intermediate, or high DLL3 expression. Genomic DNA was isolated from each sorted population, and sgRNA cassettes were PCR-amplified and deep sequenced for quantification to identify genes required for DLL3 expression (Fig. 1C). The overall performance of both screens was supported by the behavior of sgRNAs targeting *ASCL1*, which is present in both libraries and is a known activator of DLL3^41,42^, as well as sgRNAs targeting *DLL3* itself, which are present in the genome-wide library (Figs. 1D–E, Supplementary Table 2).

### Two independent CRISPR screens identify POU2F1 as a potent activator of DLL3 expression in SCLC

Both CRISPR screens independently identified POU2F1 as one of the strongest positive regulators of DLL3 expression in NCI-H209 cells (Figs. 1D–E, Supplementary Table 2). In addition to POU2F1 as a genetic outlier, the genome-wide screen also revealed enrichment for general mRNA processing factors as being needed for DLL3 expression (Figs. 1E, S1B), consistent with a broader requirement for transcriptional and post-transcriptional machinery for DLL3 expression. POU2F1 is a ubiquitously expressed member of the POU homeodomain transcription factor family, and its requirement for DLL3 expression was unexpected based on prior literature. Because POU2F1 has not been previously studied in neuroendocrine lineages or in SCLC, we focused subsequent analyses on validating its role in DLL3 regulation and assessing whether it contributes more broadly to neuroendocrine identity in SCLC.

Analysis of *POU2F1* expression across 1,699 human cancer cell lines confirmed its widespread expression (Fig. S1C, Supplementary Table 1). However, when cancer types were ranked by median *POU2F1* mRNA levels, SCLC lines exhibited the highest expression among all cancer types examined (Fig. S1C). In both cancer cell lines and human tumors, *POU2F1* expression was higher in SCLC than in other lung cancer subtypes and was relatively uniform across the molecular subtypes of SCLC (Figs. 1F–G, S1C, Supplementary Tables 1, 3)^43,44^. This pattern contrasts with *DLL3* expression, which is selectively elevated in neuroendocrine SCLC-A relative to tuft cell-like SCLC-P subtypes (Figs. 1F–G, S1D, Supplementary Tables 1, 3). Together, these findings suggest that while POU2F1 is highly expressed in SCLC, it is not sufficient on its own to dictate DLL3 expression or neuroendocrine identity.

To validate the screening results, we performed Western blot analysis following CRISPR-mediated knockout (KO) of *POU2F1* in three independent neuroendocrine SCLC cell lines (NCI-H209, NCI-H1836, and NCI-H1436). Loss of POU2F1 resulted in a reduction of DLL3 protein levels in all three models (Fig. 2A). Consistent with a transcriptional mechanism, RNA-seq analysis demonstrated a significant decrease in *DLL3* mRNA expression following POU2F1 KO (Fig. 2B, Supplementary Table 4). As a control, CRISPR-mediated inactivation of *ASCL1* similarly reduced *DLL3* mRNA and protein levels in these lines (Figs. 2A–B, Supplementary Table 4). Collectively, these data validate POU2F1 as a transcriptional activator required for DLL3 expression in multiple SCLC contexts.

**Figure 2.**
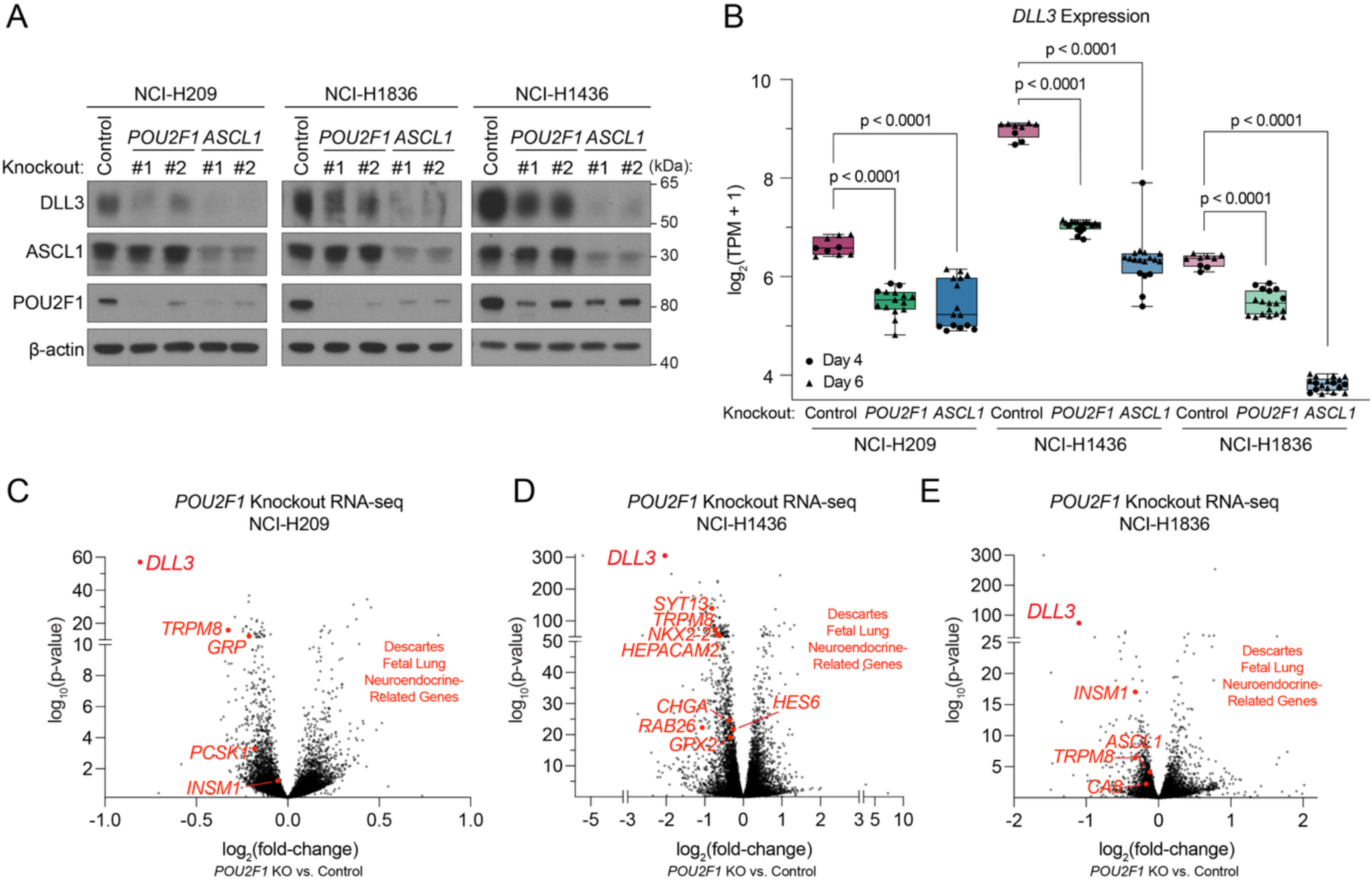
POU2F1 is required for DLL3 expression in SCLC. (A) Western blot of DLL3, ASCL1, and POU2F1. Protein lysates were collected from NCI-H209, NCI-H1836, and NCI-H1436 on day 7 following CRISPR-Cas9 knockout (KO) of the *ASCL1*, *POU2F1*, or *ROSA26* (control). Representative of 2 biological replicates. β-actin, loading control. (B–E) RNA sequencing in NCI-H209 (B–C), NCI-H1436 (B, D), and NCI-H1836 (B, E) on day 4 (B–C) or day 6 (B, D–E) following CRISPR-Cas9 knockout (KO) of *POU2F1, ASCL1,* or *ROSA26* (control). (B) *DLL3* expression in transcripts per million (TPM). Each point = 1 biological replicate. The day post-KO is indicated by the shape. Boxes represent the mean ± interquartile range (IQR). Error bars extend to the minimum and maximum values. Statistical significance was evaluated using two-sided Mann–Whitney tests with no multiple comparison correction. (C–E) Volcano plots of differentially expressed genes following *POU2F1* KO. Fold change and significance calculated by DESeq2. n = 6 biological replicates per timepoint, per cell line. *DLL3* and additional genes related to fetal lung neuroendocrine identity (DESCARTES database; Cao et al. 2021) are labeled.

### POU2F1 supports the neuroendocrine transcriptional program in SCLC

We next examined the broader transcriptional consequences of POU2F1 inactivation in SCLC. Comparative analysis of RNA-seq datasets revealed that *DLL3* was among the most significantly and strongly downregulated genes following *POU2F1* KO across all three SCLC-A cell lines (Figs. 2B–E, Supplementary Table 4). Additional genes exhibiting strong POU2F1 dependence included canonical markers of neuroendocrine identity such as *CHGA*, *INSM1*, *GRP*, and *TRPM8* (Figs. 2C–E, Supplementary Table 4)^45^. To assess whether these changes reflected a coordinated loss of neuroendocrine identity, we performed Gene Set Enrichment Analysis^46^ using gene signatures associated with ASCL1-positive neuroendocrine SCLC (SCLC-A) and with normal human fetal lung neuroendocrine cells derived from the Developmental Single Cell Atlas of Gene Regulation and Expression (DESCARTES) database^47^. Both signatures showed a significant reduction following POU2F1 KO (Figs. 3A–F). Unbiased Gene Ontology analysis further identified “Lung Neuroendocrine genes” as the most significantly enriched category among genes downregulated upon POU2F1 KO (Figs. 3G–I)^48^. Parallel analyses of *ASCL1*-KO transcriptomes in these same cell line models yielded similar results (Figs. S2A–L, Supplementary Table 4). Together, these findings indicate that POU2F1 and ASCL1 each perform essential roles in sustaining the neuroendocrine transcriptional program of SCLC cells.

**Figure 3.**
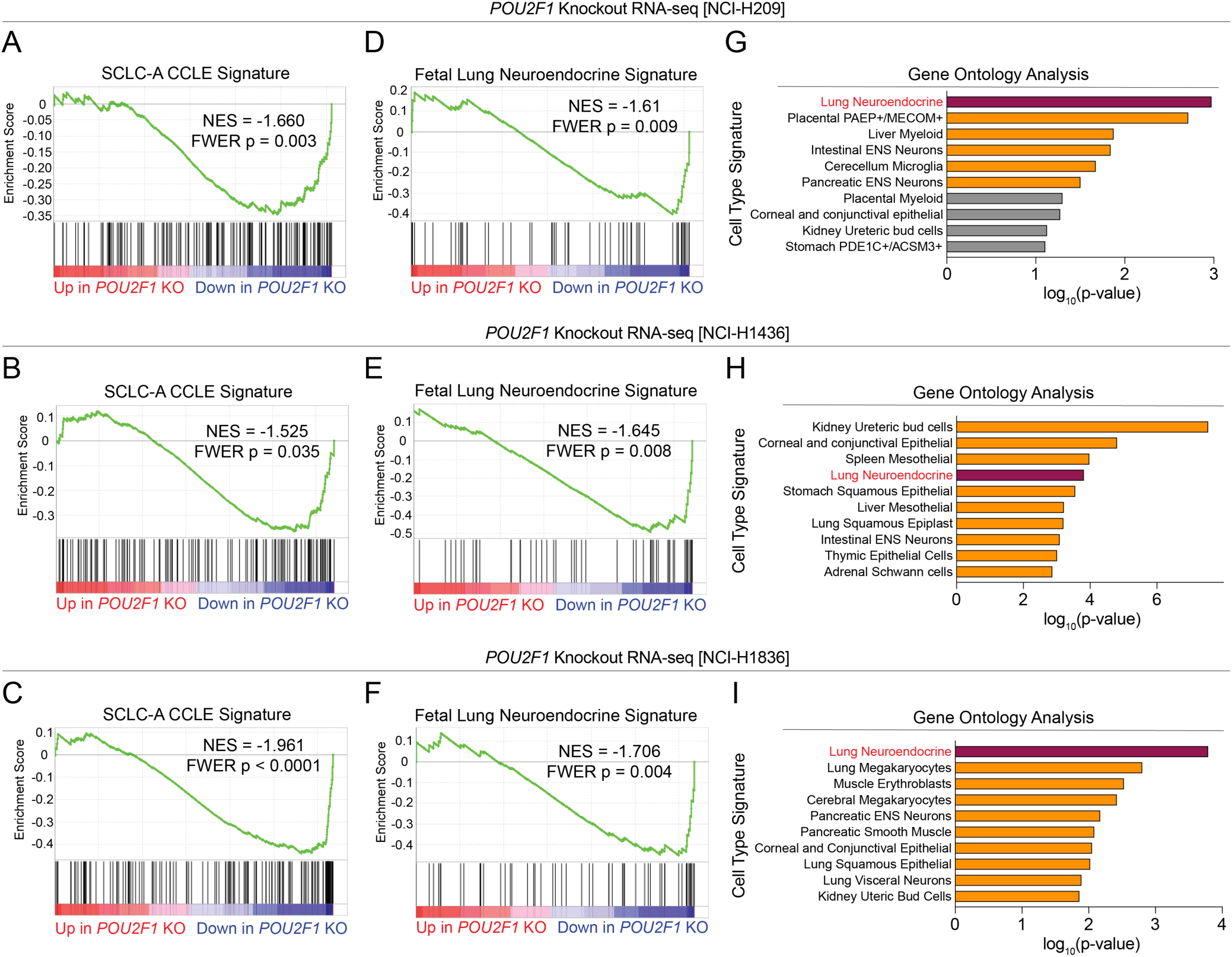
POU2F1 activates a neuroendocrine identity gene expression program in three independent SCLC-A cell lines. (A–I) RNA sequencing in NCI-H209 (A,D,G), NCI-H1436 (B,E,H), and NCI-H1836 (C,F,I) on day 4 (A,D,G) or day 6 (B,C,E,F,H,I) following CRISPR-Cas9 knockout (KO) of *POU2F1*, or *ROSA26* (control). Fold change and significance calculated by DESeq2. n = 6 biological replicates per timepoint, per cell line. (A–F) Gene set enrichment analysis (GSEA) of differentially expressed genes following *POU2F1* KO. Significance calculated by GSEA. NES = Normalized Enrichment score. FWER = Family-wise Error Rate. (G–I) Enrichr Gene ontology analysis of top 250 downregulated genes following *POU2F1* KO. Top 10 DESCARTES Cell Type signatures are ranked ordered by their significance (p-value) and the most significant terms (p < 0.05) are highlighted.

### POU2F1 and ASCL1 both occupy cis-regulatory elements at neuroendocrine genes

We next asked whether POU2F1 directly regulates *DLL3* and other neuroendocrine genes and whether its activity is integrated with ASCL1 at the level of chromatin occupancy. Using two independent antibodies validated for chromatin immunoprecipitation, we performed POU2F1 ChIP-seq in NCI-H209 cells. Although the total number of high-confidence POU2F1 peaks was relatively modest (fewer than 700), the datasets showed concordance between antibodies and robust enrichment of the known POU2F1 DNA recognition motif (Figs. 4A–C, Supplementary Table 5). Motif enrichment analysis^49^ also identified the ASCL1 recognition sequence as significantly enriched within POU2F1-occupied regions (Fig. 4C, Supplementary Table 5). This observation prompted us to perform ASCL1 ChIP-seq in the same cell line, which confirmed co-occupancy of POU2F1 and ASCL1 at a subset of cis-regulatory elements (Figs. 4B, D, S3A–C). Notably, both transcription factors occupied the *DLL3* promoter (Fig. 4D). Reciprocal motif analysis of ASCL1 peaks revealed enrichment of the POU2F1 recognition motif (Fig. S3D, Supplementary Table 5). Consistent with this observation, the *DLL3* promoter contains tandem POU2F1 and ASCL1 motifs (Fig. 4E). A similar pattern of occupancy was observed at a regulatory element located approximately 6 kilobases upstream of the *INSM1* gene, again in association with tandem POU2F1 and ASCL1 motifs (Figs. 4D–E).

**Figure 4.**
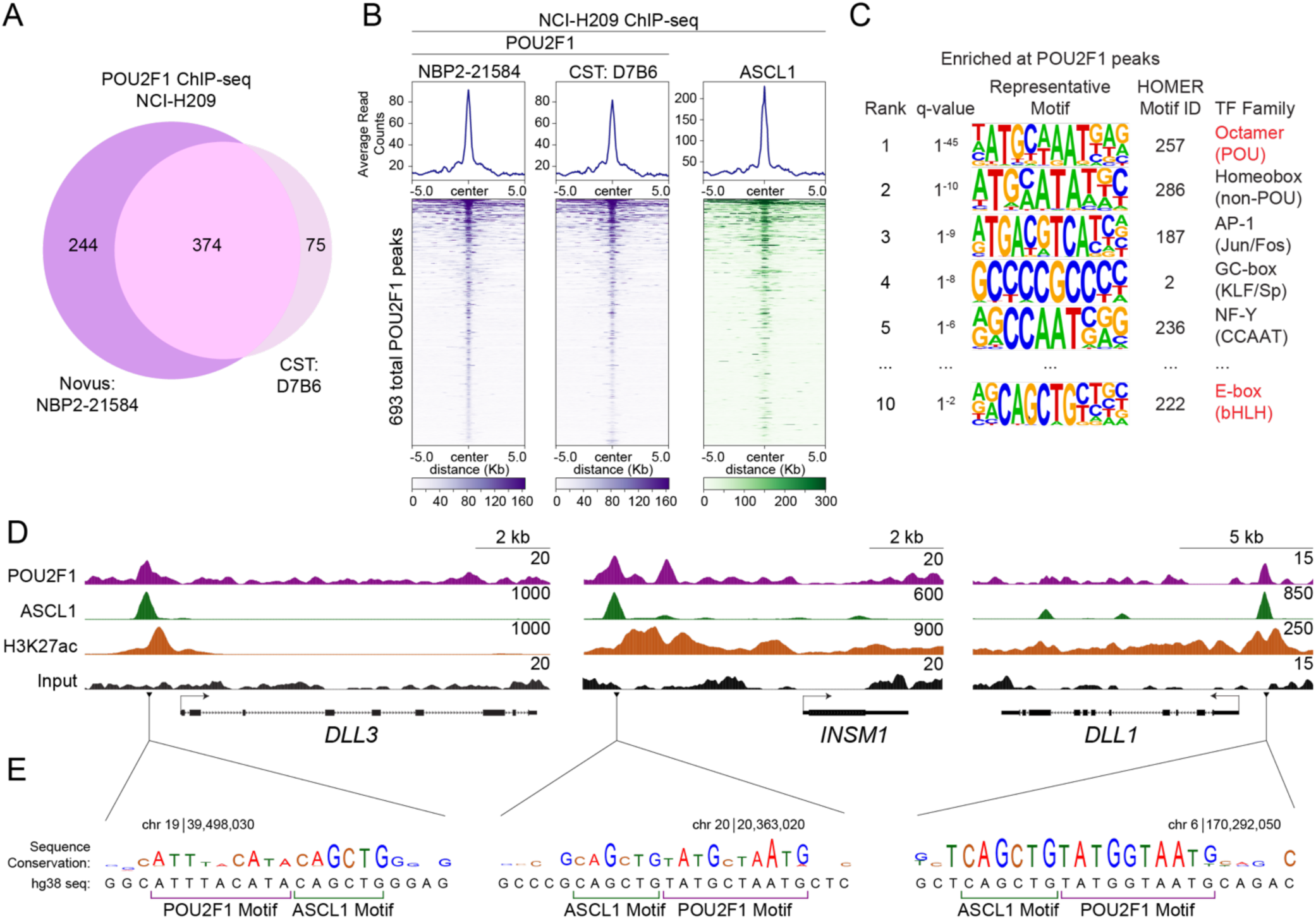
POU2F1 and ASCL1 occupy chromatin at tandem POU2F1–ASCL1 motifs. (A) Venn diagram of 693 total POU2F1 peaks (MACS2 q < 0.01) in NCI-H209 cells, including IP replicates using two independent antibodies (denoted). (B) Heatmaps for POU2F1 and ASCL1 ChIP-seq at all POU2F1 peaks. Rows = 10kb genomic regions centered on a POU2F1 peak summit. Rows are ordered by POU2F1 signal, and this ordering is applied to both heatmaps. Metagene plots show the average signal for each factor across all peaks, plotted above each heatmap. (C) HOMER motif enrichment analysis of 693 called POU2F1 peaks (MACS2 q < 0.01). The top 5 transcription factor family motifs were selected. (D) ChIP-seq tracks in NCI-H209 showing ASCL1, POU2F1, H3K27ac, and input enrichment at representative loci, visualized in UCSC genome browser. Matched, scaled track heights are listed (right). (E) hg38 sequence (bottom) and sequence conservation across 100 vertebrates (top) at the locus of ASCL1/POU2F1 peak summits adjacent to *DLL3*, *INSM1*, and *DLL1*. Base height indicates conservation across 100 vertebrates in UCSC genome browser. Genomic location (above) and locations of POU2F1 and ASCL1 motifs (below) are labeled.

### Tandem POU2F1–ASCL1 motifs contribute to a neuroendocrine identity cis-regulatory code

These findings led us to hypothesize that tandem POU2F1–ASCL1 motifs represent a feature of cis-regulatory elements that control neuroendocrine identity. To test this idea, we constructed a custom position weight matrix for tandem POU2F1–ASCL1 motifs and scanned the human genome (hg38) for occurrences of this composite element using the FIMO algorithm (Fig. 5A)^50^. This analysis identified 882 instances of tandem motifs in the human genome (Supplementary Table 6). Most tandem motifs were located at distal intergenic or intragenic regions rather than promoters, with *DLL3* representing a notable exception (Fig. S4A). Using the HOMER algorithm, we assigned each tandem motif to its most likely target gene, yielding a set of 227 genes within 20 kilobases of a tandem POU2F1–ASCL1 motif^49^. Remarkably, a Gene Ontology analysis of this gene set revealed “Lung Neuroendocrine” as the top enriched category associated with the presence of a nearby tandem motif (Fig. 5B)^48^. Importantly, this enrichment was substantially stronger for the tandem motif than for either the POU2F1 or ASCL1 motif alone and this effect was orientation-dependent, indicating a unique regulatory association of the composite element (Figs. 5B–D, S4B, Supplementary Table 6).

**Figure 5.**
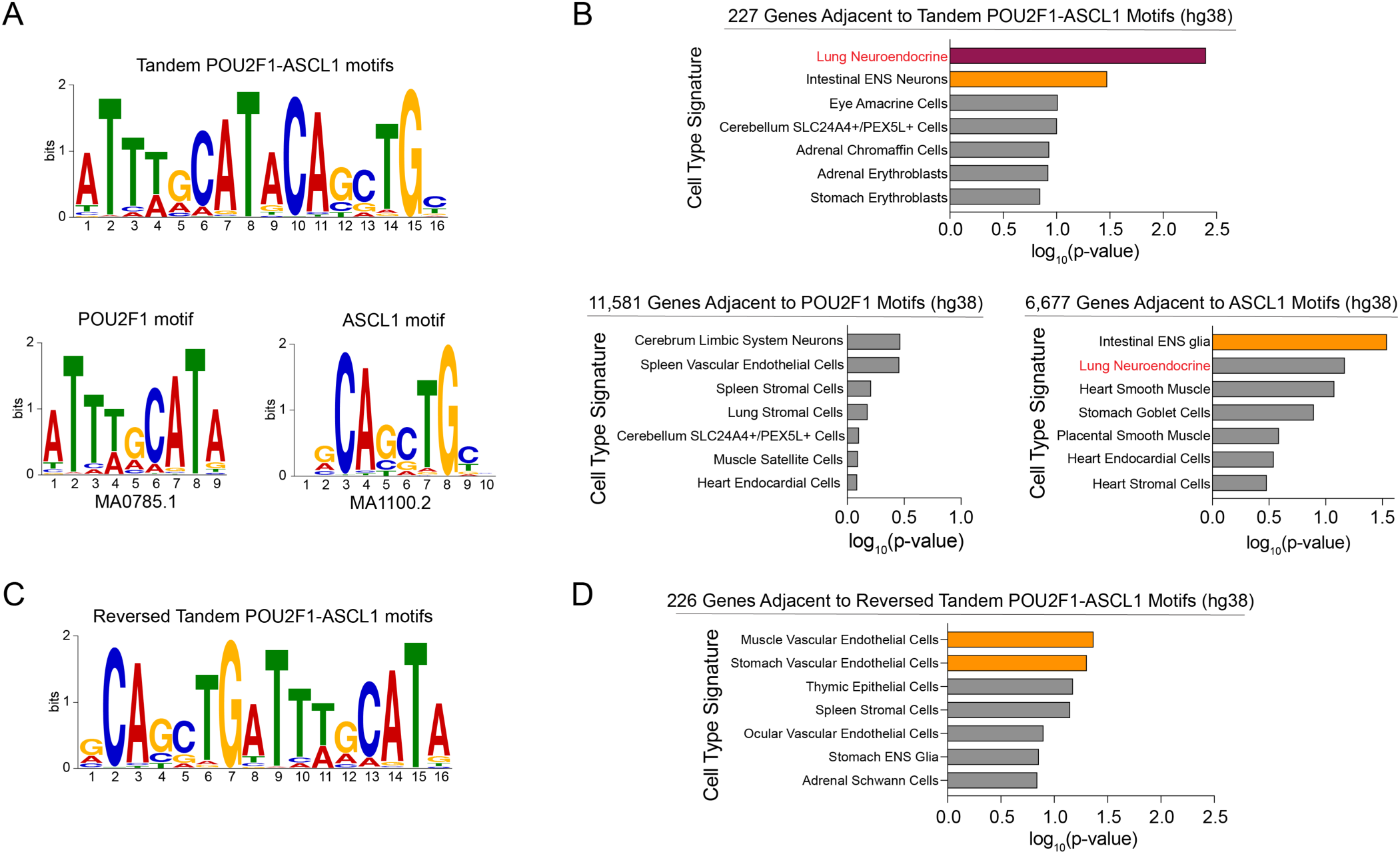
Tandem POU2F1–ASCL1 motifs represent a lung neuroendocrine identity-defining cis-regulatory grammar. (A, C) Schematic of the ASCL1, POU2F1, tandem POU2F1–ASCL1, and reversed tandem POU2F1–ASCL1 motifs. Position weight matrices generated using Tomtom (MEME suite). Height (bits) represents the conservation of the annotated base pair at that position. Jaspar Motif IDs are labeled below. (B, D) Enrichr Gene ontology analysis of annotated genes within 20kb of a tandem POU2F1–ASCL1 (p < 1×10^7^), reversed tandem POU2F1–ASCL1 (p < 1×10^7^), POU2F1 (p < 1×10^5^), or ASCL1 (p < 1×10^5^) motif (hg38). Top 7 DESCARTES Cell Type signatures are ranked ordered by their significance (p-value) and the significant terms (p < 0.05) are highlighted.

We next hypothesized that a tandem motif architecture might permit DNA-dependent interactions between POU2F1 and ASCL1. Because ASCL1 functions as a heterodimer with E proteins such as TCF12^51^, we used multiple AlphaFold 3 predictions to generate structural models of the ASCL1–TCF12 heterodimer in complex with POU2F1 and a 16 bp dsDNA fragment containing tandem POU2F1 and ASCL1 motifs (5’-**ATTTGCATA***CAGCTG*T-3’ and complement, Fig. 6A)^52^. Across multiple predictions, the structured regions of all three proteins were positioned with high confidence relative to the DNA, whereas the relative placement of the proteins with respect to one another showed greater variability (Figs. 6A–C, S5A–C). One recurrent higher-confidence feature was a predicted interaction between a charged cluster at the N-terminus of the TCF12 bHLH domain and a basic patch in POU2F1 encompassing residues 377–382, located adjacent to the POU2F1 DNA motif (Figs. 6C–D, S5A–B). In addition, a short region of ASCL1 immediately N-terminal to its bHLH domain was predicted to contact POU2F1 in two distinct conformations, albeit in lower confidence than the charged interaction (Figs. 6C, 6E, S5A–B). Both the overall model confidence and the relative confidence in protein positioning were markedly reduced when the tandem motif orientation was reversed (5’-*CAGCTG*T**ATTTGCATA**-3’ and complement) or when DNA was omitted from the prediction, consistent with a DNA-stabilized interaction between POU2F1 and the ASCL1–TCF12 complex (Figs. 6B–C, S5A–D). Together, these findings are consistent with a model in which tandem POU2F1 and ASCL1 motifs facilitate DNA-dependent interactions between POU2F1 and the ASCL1–TCF12 complex.

**Figure 6.**
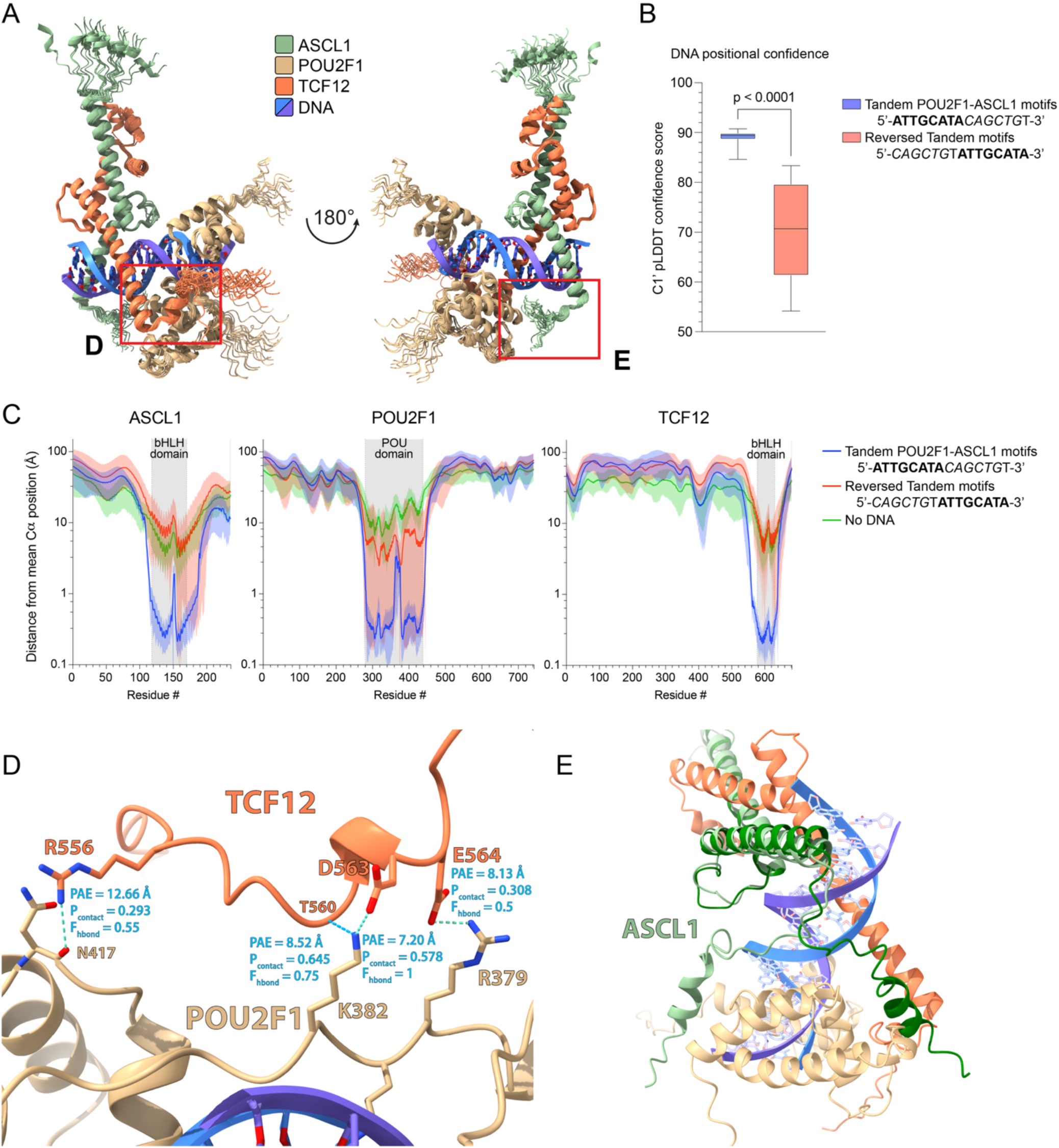
AlphaFold 3 supports an orientation-specific tandem POU2F1-ASCL1 motif. (A) Cartoon schematic of twenty (four starting seeds, five models each) aligned AlphaFold 3 predictions of the tandem motif bound to the ASCL1–POU2F1–TCF12 complex. (B) The C1’ carbon is the carbon atom within the deoxyribose sugar of DNA that forms the covalent bond between the nitrogenous base and the sugar backbone. Higher pLDDT scores at DNA nucleotide positions indicate greater model confidence in the predicted placement of the DNA, reflective of more confident positioning of the DNA within the complex, and suggestive of stabilization of the DNA structure by bound proteins. Confidence scores (pLDDT) in the placement of the DNA nucleotides (N=32 each) in AlphaFold 3 models containing the ASCL1–POU2F1 tandem motif or a “reversed” motif are plotted. Each data point represents the mean pLDDT score at a given nucleotide position in the DNA sequence. Boxes represent the mean ± interquartile range (IQR). Error bars extend to the minimum and maximum values. Statistical significance was evaluated using Welch’s unpaired t-test. **Bold letters** were used to represent the POU2F1 binding motif, while *italics* indicate the ASCL1 binding motif. (C) The Cα carbon is the central backbone carbon of each amino acid, bonded to the amino, carboxyl, and side chain groups. Smaller distances from the mean aligned position indicate greater model-to-model consistency across AlphaFold 3 predictions, suggestive of a more stably predicted local protein structure. The distance of each residue’s Cα atom from the mean aligned position of predictions containing the tandem motif sequence, a “reversed” tandem motif, or models that did not include DNA are plotted. Central line represents the mean distance from the aligned Cα, with the standard deviation of the mean residue distance surrounding it. (D) The predicted POU2F1–TCF12 interface as seen in a representative AlphaFold 3 prediction. PAE and contact probability values are represented by the mean of those values over all twenty models in both directions. F_hbond_ represents the fraction of AlphaFold 3 predictions in which a hydrogen bond was projected to occur using relaxed constraints in ChimeraX. (E) The N-terminal region of ASCL1, as shown in two representative AlphaFold 3 predictions, fell into one of two distinct conformations in most predictions.

Finally, we assessed whether genes linked to tandem POU2F1–ASCL1 motifs were functionally dependent on these transcription factors. Gene Set Enrichment Analysis demonstrated significant downregulation of tandem-associated genes following *POU2F1* KO across all three SCLC cell lines (Fig. 7). This pattern was not recapitulated when examining genes associated with POU2F1 motifs alone (Fig. S6A). Similar results were observed following *ASCL1* inactivation in the same cell lines (Figs. S6B–C). Taken together, these data support a model in which tandem POU2F1–ASCL1 motifs serve as critical cis-regulatory elements that integrate the activities of both transcription factors to maintain neuroendocrine identity in SCLC, and potentially in normal neuroendocrine cellular contexts.

**Figure 7.**
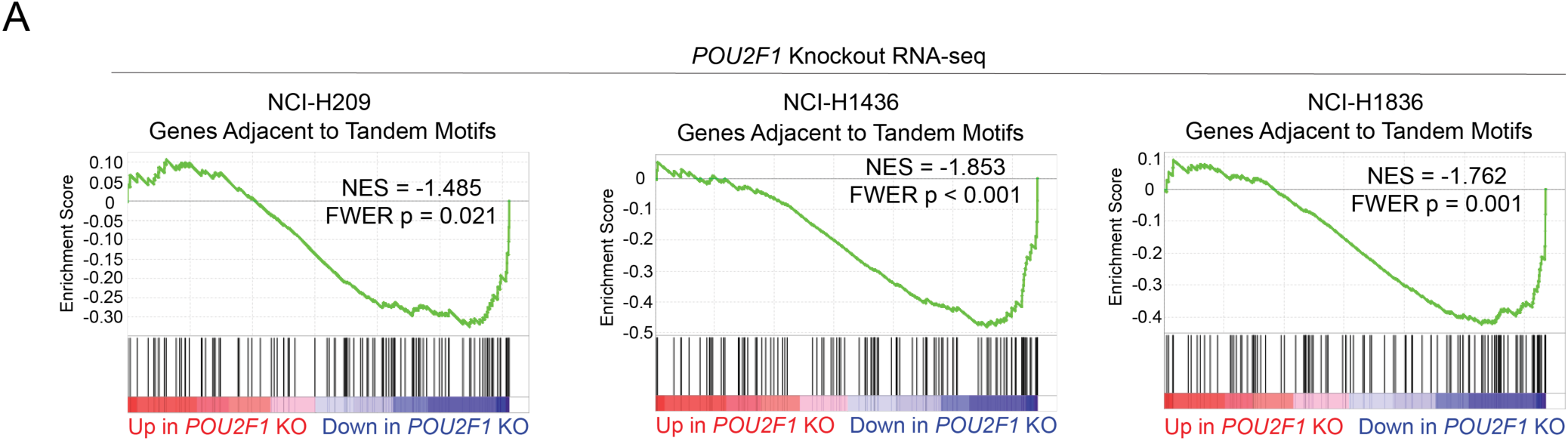
Tandem POU2F1–ASCL1 motifs correlate with POU2F1-activated gene expression in small cell lung cancer cell lines. (A) RNA sequencing in NCI-H209, NCI-H1436, and NCI-H1836 following CRISPR-Cas9 knockout (KO) of *POU2F1*, or *ROSA26* (control). Fold change and significance calculated by DESeq2. n = 6 biological replicates per timepoint, per cell line. Gene set enrichment analysis (GSEA) of differentially expressed genes following *POU2F1* KO. NES = Normalized Enrichment score. FWER = Family-wise Error Rate.

## Discussion

This study indicates that DLL3 expression in SCLC is governed by a defined cis-regulatory logic rather than serving as a passive marker of neuroendocrine identity. While DLL3 has been widely used as a biomarker and therapeutic target, its regulation has largely been inferred indirectly through associations with ASCL1-positive tumors and suppressed Notch signaling^9,41,53^. By applying unbiased CRISPR screening approaches, we directly map the transcriptional requirements that sustain DLL3 expression and demonstrate that high-level DLL3 transcription requires coordinated activity of multiple transcription factors. These findings position DLL3 as an encoded output of a lineage-specific transcriptional program, with implications for understanding both its biological role and its behavior under therapeutic pressure.

A key insight from this work is that a ubiquitously expressed transcription factor, POU2F1, acquires a lineage-restricted function in SCLC through cooperative interactions with ASCL1 within a specific cis-regulatory architecture. POU2F1 has traditionally been viewed as a broadly acting transcriptional regulator, rather than as a determinant of cell identity^30^. Our findings add to a growing body of evidence challenging this view by showing that POU2F1 becomes essential for neuroendocrine gene expression in SCLC when paired with ASCL1 at tandem DNA motifs^37–39^. One potential explanation for this context specificity is the pioneer factor activity of ASCL1. In multiple developmental and reprogramming systems, ASCL1 can bind nucleosome-occluded chromatin and induce chromosome remodeling, subsequently recruiting additional transcriptional regulators^54–56^. It is possible that, in SCLC, ASCL1 functions as the primary lineage-defining factor that establishes accessible chromatin at neuroendocrine cis-regulatory elements, in turn enabling binding of POU2F1 to tandem POU2F1–ASCL1 motifs, which leads to functional cooperativity via a DNA-dependent interaction between these two transcription factors. Consistent with this model, ASCL1 chromatin binding during induced fibroblast-to-neuronal reprogramming precedes recruitment of BRN2, a POU family transcription factor, which cannot productively access chromatin alone^57^. Following ASCL1 binding, BRN2 effectively binds neuronal cis-regulatory elements and activates transcription to support reprogramming. This observation supports a similar hierarchical model of chromatin engagement during SCLC progression in which ASCL1 initiates the global regulatory landscape, while POU2F1 reinforces transcriptional activation at a defined subset of neuroendocrine-specific loci.

These findings also have implications for understanding lineage plasticity in SCLC and how tumors might transition between neuroendocrine and non-neuroendocrine states. Prior work has shown that tuft cell-like SCLC is driven by a distinct transcriptional circuitry centered on POU2F3, often in conjunction with ASCL2, a paralog of ASCL1^22,24^. The parallels between POU2F1–ASCL1 cooperation in neuroendocrine SCLC and POU2F3–ASCL2 function in tuft cell-like carcinomas suggest a modular transcriptional system in which different POU and ASCL family members pair to specify alternative lineage programs. Such pairing could provide a mechanism for lineage switching, whereby changes in expression or activity of one component enable engagement of an alternative cis-regulatory code. Future investigations will define the molecular determinants that govern selective pairings between POU and ASCL family members and will determine whether lineage transitions reflect modified transcription factor paralog pairings, competition for access to shared cis-regulatory elements, or the activation of distinct enhancer networks. A more comprehensive characterization of how alternative POU–ASCL modules are assembled and reprogrammed during cancer progression and upon therapeutic pressure may reveal vulnerabilities that can be exploited to constrain lineage plasticity or to redirect tumor identity.

An improved understanding of DLL3 transcriptional regulation has direct relevance for DLL3-directed therapeutic strategies in SCLC. Clinical efforts targeting DLL3 assume relative stability of antigen expression, yet emerging evidence indicates that SCLC tumors can alter lineage state in response to treatment^58^. Our findings suggest that DLL3 expression is tightly linked to a specific transcriptional configuration, raising the possibility that disruption of this regulatory logic could lead to antigen loss and therapeutic resistance. Conversely, persistence of the POU2F1–ASCL1 regulatory axis may help explain sustained DLL3 expression in a subset of tumors. These considerations highlight the importance of transcriptional mechanisms in determining the durability of tumor-selective antigens.

## Methods

### Institutional approval

This study complies with all relevant ethical regulations, and all protocols were approved by the Cold Spring Harbor Laboratory Institutional Biosafety Committee (IBC).

### Cancer cell lines and tissue culture

The following cell lines used in this study were obtained from ATCC: HEK293T (Cat# CRL-3216; RRID:CVCL_0063), NCI-H209 (male, Cat# HTB-172; RRID:CVCL_1525), NCI-H1836 (male, Cat#: CRL-5898; RRID:CVCL_1498), and NCI-H1436 (male, Cat# CRL-5871; RRID:CVCL_1471).

HEK293T cells were cultured in DMEM supplemented with 10% FBS. NCI-H209, NCI-H1436, and NCI-H1836 cells were cultured in HITES media supplemented with 5% FBS. HITES media is composed of DMEM:F12 base medium supplemented with 0.005 mg/mL insulin, 0.01 mg/mL transferrin, 30 nM sodium selenite, 10 nM hydrocortisone, 10 nM β-estradiol, and 4.5 mM L-glutamine. 1% Penicillin–streptomycin was added to all media. Cell lines were purchased from commercial vendors and their identity was validated by STR analysis. Cell lines were regularly tested for *Mycoplasma* contamination. All antibiotic concentrations used to select gene cassettes were empirically titrated in each cell line to achieve maximum selection with minimum toxicity.

### Lentiviral Production and Infection

Lentivirus was produced in HEK293T cells transfected with target plasmids and packaging plasmids (VSV-G and psPAX2) using polyethyleneimine. Transfection media was replaced with fresh HITES media supplemented with 5% FBS 6 hours after transfection, and lentivirus-containing supernatant was collected 24, 48, and 72 hours following transfection. All three collections were pooled and filtered using a 0.45 µm PES filter. For lentiviral infections, cell suspensions were exposed to lentiviral-containing supernatant supplemented with polybrene to a final concentration of 4 µg/mL and spun at 600 RCF for 30 minutes. Lentiviral media was replaced with fresh media after 24 hours.

### Intracellular FACS-based CRISPR Screens

The lentiGuide-Puro backbone (Addgene #52963) containing the genome-wide Brunello^40^ (Addgene #73178) sgRNA library was first modified prior to library cloning to replace the puromycin-resistance cassette with a GFP-P2A-hygromycinR cassette. After empirical determination of the suitable lentiviral titer, ∼5 × 10^7^ (transcription factor-focused screen) or 7.5 × 10^8^ (genome-wide screen) Cas9-expressing cells were infected with human DNA Binding Domain-Focused^22^ (Addgene #123334) or genome-wide Brunello sgRNA library-encoding lentiviral suspension for a 20–30% infection efficiency. Media was replaced with HITES media at 48h. For the genome-wide screen, hygromycin B was added for 72 hours to select for transduced cells. 8 days post infection, cells were collected, treated with Accumax^TM^ (ThermoFisher) to dissociate cell clumps, washed in PBS, and then fixed in −20°C methanol at ≤ 10 × 10^6^ cells/mL under gentle vortexing. Cells were stored in methanol at −20°C for at least 2 days and up to 1 month. One day before sorting, cells were pelleted, washed 1x in FACS buffer (1% (w/v) ultrapure BSA, 0.5% (w/v) sodium azide, and 1 mM EDTA in magnesium and calcium-free PBS), and incubated overnight in 1:200 primary antibody (DLL3) in FACS buffer at 10 × 10^6^ cells/mL rotating at 4°C. The next day, cells were pelleted, washed twice with FACS buffer, and incubated for 2 hours in 1:500 secondary antibody (AlexaFluor647-conjugated anti-rabbit) in FACS buffer at 10 × 10^6^ cells/mL rotating at 4°C protected from light. After washing 2x in FACS buffer, cells were resuspended in FACS buffer at 10 × 10^6^ cells/mL and sorted. Stained cells were sorted using a BD FACS Aria II cell sorter. The total number of cells sorted per screen was a minimum of 2500x the size of the sgRNA library. Cells were sorted into three different populations, with approximately 20% of the cells sorted into the DLL3^low^ bin, 70% of cells sorted into the DLL3^bulk^ bin, and 10% of the cells being sorted into the DLL3^high^ bin. Cell pellets were then processed for DNA extraction and library preparation as described below. Custom sequencing primers were added for each respective cell population. All sequencing data from FACS-based screens was analyzed with MAGeCK^59^ v0.5.9.3 using the MLE option. For the transcription factor-focused DLL3 marker-based screen, the DLL3^high^ gate was insufficiently diverse for downstream analysis, so the DLL3^low^ and DLL3^bulk^ gates were used to analyze this screen.

### DNA extraction and sgRNA sequencing for CRISPR screens

After pooling and pelleting of sorted cells, cells were resuspended in DNA extraction buffer (10 mM Tris-HCl pH 8.0, 150 mM NaCl, and 10 mM EDTA) at a density of ≤ 5 × 10^7^ cells/mL. SDS and Proteinase K were added to final concentrations of 0.1% and 0.2 mg/mL, respectively. The mixture was incubated for 48 hours at 56°C, after which DNA was purified by phenol extraction. Equilibrated phenol was added 1:1 to the lysis mixture, mixed well, and centrifuged for 10 minutes at 20,000 RCF. Supernatant was carefully removed, and another phenol purification round was performed. DNA was then precipitated by adding 3 volumes of isopropanol and NaOAc pH 5.2 to a final concentration of 75 mM and incubating overnight at −20°C. DNA was pelleted at 20,000 RCF for 1 hour, washed in 70% ethanol, and air-dried until translucent. After resuspension in sterile, ultrapure water, DNA was assessed for quality by NanoDrop before proceeding to library prep. sgRNAs were directly amplified from genomic DNA by one step PCR using NEBNext^®^ Ultra^TM^ II Q5^®^ Master Mix (NEB). Each PCR reaction was performed with 10 μg of genomic DNA in 100 µL final volume. Titrations of amplification cycles were performed for LRG or LentiCRISPRv2 (95°C, 1 min; n cycles [95°C, 30s; 53°C, 30s; 72°C, 30s]; 72°C, 10 min) sgRNA cassettes. All PCR reactions for each sample were pooled and 400 µL were taken for double-sided AMPure bead cleanup (0.65x + 1x bead volume to preserve PCR amplicons (∼192 and ∼274 bp, respectively)). Amplicons were sequenced using an Illumina NextSeq with 50% spike-in or pooled with high-diversity libraries (Cold Spring Harbor Genome Center, Woodbury, NY 11797).

### RNA extraction, RT-qPCR, and RNA-sequencing

Total RNA was extracted using TRIzol reagent following manufacturer’s instructions. For RNA-seq experiments following CRISPR-based targeting of *POU2F1*, *ASCL1*, or controls, cells stably expressing Cas9 were infected with control or target sgRNAs in an LRG2.1_Blast vector (Addgene #65656) to >90% GFP positivity. RNA was collected at the indicated timepoints (4 days or 6 days following lentiviral transduction).

RNA-sequencing libraries were constructed using the TruSeq sample Prep Kit V2 (Illumina) following manufacturer’s instructions. Briefly, 2 µg of extracted, purified RNA was poly-A selected and fragmented with fragmentation enzyme mix. cDNA was synthesized with SuperScript^TM^ II reverse transcriptase, followed by end repair, A-tailing, single-end indexed adaptor ligation, and PCR amplification. RNA-sequencing libraries were single-end sequenced for 76 bp using an Illumina NextSeq platform (Cold Spring Harbor Genome Center, Woodbury, NY 11797).

### ChIP and ChIP-seq library construction

For each ChIP, cells were collected, washed 2x with room temperature PBS, resuspended in Accumax^TM^ (ThermoFisher) at 10×10^6^ cells/mL and rotated at room temperature for 2 hours. Cells were then washed 2x with room temperature PBS and resuspended at 5-10 × 10^6^ cells/mL in room temperature PBS. Cell suspensions were crosslinked in 1% formaldehyde at room temperature for 15 minutes, followed by the addition of glycine to quench the reaction at a final concentration of 0.125 M. After two ice-cold PBS washes, cells were resuspended in cell lysis buffer (10 mM Tris pH 8.0, 10 mM NaCl, 0.2% NP-40) at 10×10^6^ cells/mL and incubated on ice for 15 minutes. After spinning down, supernatant was removed and nuclei were resuspended in nuclear lysis buffer (50 mM Tris pH 8.0, 10 mM EDTA pH 8.0, 1% SDS) at 20×10^6^ cells/mL and sonicated in 15 mL tubes using a Bioruptor^®^ Pico water bath sonicator (13 × 30s on/off cycles, 1 mL per tube) to achieve an average chromatin size distribution of 200–500 base pairs. Each 1 mL of sonicated chromatin from 20 million cells was diluted with 7 mL of IP-Dilution buffer (20 mM Tris pH 8.0, 2 mM EDTA pH 8.0, 150 mM NaCl, 1% Triton X-100, 0.01% SDS), and 200 µL of the sample was saved for input. Chromatin from 20 million (H3K27ac) or 100 million (POU2F1, ASCL1) cells was incubated with 4 µg (H3K27ac) or 10 µg (POU2F1, ASCL1) of the appropriate antibody and 25-100 µL of magnetic protein-A (rabbit) or protein-G (mouse) beads at 4°C overnight. Beads were then pooled for each respective IP and washed once with 1 mL IP-wash buffer 1 (20 mM Tris pH 8.0, 2 mM EDTA pH 8.0, 50 mM NaCl, 1% Triton X-100, 0.1% SDS), twice with 1 mL High-salt buffer (20 mM Tris pH 8.0, 2 mM EDTA pH 8.0, 500 mM NaCl, 1% Triton X-100, 0.01% SDS), once with IP-wash buffer 2 (10 mM Tris pH 8.0, 1 mM EDTA pH 8.0, 250 mM LiCl, 1% NP-40, 1% sodium deoxycholate), and twice with 1 mL TE buffer (10 mM Tris-Cl, 1 mM EDTA, pH 8.0). Chromatin was eluted from beads and reverse-crosslinked in 200 µL nuclear lysis buffer supplemented with 12 µL NaCl and 1 µg/mL RNase A by shaking at 800 rpm for ≥ 4 hours at 65°C. Supernatant was isolated from magnetic beads and protein digestion was performed by adding 4 µg/mL of Proteinase K and incubating the mixture for 2 hours at 56°C. NaOAc pH 5.2 was added to a final concentration of 75 mM and the DNA to be used for library prep was purified using the QIAGEN PCR purification kit and following manufacturer’s instructions.

Each ChIP-seq library was constructed using the Illumina TruSeq ChIP Sample Prep kit following manufacturer’s instructions. Briefly, ChIP DNA was end repaired, A-tailed, and ligated to Illumina-compatible single-indexed adaptors. 15 PCR cycles were used for final library amplification. After amplification, the library was purified 2x with 1x AMPure XP beads and analyzed on a Bioanalyzer using a high sensitivity DNA chip (Agilent). Library DNA concentrations were quantified using an Invitrogen Qubit 4 Fluorometer using the 1x High-Sensitivity dsDNA assay. ChIP-seq libraries were single-end sequenced for 76 or 100 bp at a sequencing depth of ≥ 50 million raw reads (H3K27ac or input) or ≥ 30 million raw reads (POU2F1, ASCL1) per sample using an Illumina NextSeq platform (Cold Spring Harbor Genome Center, Woodbury, NY 11797).

### General computational, data, and statistical analyses

All sequencing data was analyzed using the CSHL High-Performance Computing system (HPC). Packages used to analyze next-generation sequencing data were installed in independent Anaconda environments to minimize dependency conflicts. Downstream analyses were performed using Python 3 in JupyterLab notebooks or in RStudio. Mann–Whitney U tests and Fisher’s Exact statistical tests were performed using R or PRISM v10. Cancer Cell Line Encyclopedia (CCLE) data were downloaded from the Depmap portal (25Q3). Human small cell lung cancer gene expression data was reanalyzed from George et al. (2015).

### RNA-sequencing analysis

Single-end 76 bp raw sequencing reads were pseudo-aligned to the hg38 genome using Kallisto^60^ with bootstrap 100. Low-abundance transcripts were removed, and variance-stabilized normalized transcripts and differential expression were calculated using DESeq2^61^. Transcripts per million (TPM) were calculated from aligned, mapped reads normalized for gene length using tximport.

Gene set enrichment analysis (GSEA) was conducted using GSEA_4.3.3^46^. The Descartes Fetal Lung Neuroendocrine GSEA Signature was downloaded from the Molecular Signatures Database (MSigDB)^47^. GSEA was performed using rank-ordered log_2_(Fold Change) values and the GSEA “Preranked” option. Gene sets are included as Supplementary Table 7. Gene Ontology analysis was conducted using Enrichr^48^ (Ma’ayan Lab) using the top 250 significantly downregulated genes (RNA-sequencing analysis), or all genes adjacent to each respective set of peaks (ChIP-sequencing analysis and motif analysis).

### ChIP-sequencing analysis

Single-end 76 base pair sequencing reads were mapped to the hg38 genome using Bowtie2^62^ with default settings. MACS2^63^ v2.2.9.1 was used to call peaks using no input genomic DNA control. Annotation and motif analysis of ChIP-seq peaks was performed using HOMER^49^ v5.11 with default settings. To visualize genomic tracks, bigWig files were generated from sorted, indexed BAM files using deepTools^64^ v3.5.2 bamCoverage function. Reads from single-end sequencing were extended based on sonication fragment size for heatmap generation (300 base pairs).

To define BED files of peaks and peak overlaps, MACS2 output narrowPeak or broadPeak files were merged using bedtools^65^ v2.30.0 intersect tools. Regions of high artifactual mapping to chromatin, “blacklisted” regions^66^, were removed from BED files using bedtools intersect prior to each analysis. Heatmaps and average chromatin occupancy metaplots were generated using computeMatrix and plotHeatmap functions of deepTools, taking bigWig and BED files as input.

### Motif Analysis

#### Motif analysis of ChIP-seq peaks was performed using HOMER^49^ v5.11 with default settings

The tandem POU2F1–ASCL1 motif was generated manually by combining position weight matrix (PWM) values for the *Homo sapiens* POU2F1 (MA0785.1) and ASCL1 (MA1100.2) motifs. The tandem POU2F1–ASCL1 motif PWM graphic was generated using the Tomtom^67^ motif comparison tool (v. 5.5.9) in the Multiple Em for Motif Elicitation (MEME) suite.

The Find Individual Motif Occurrences (FIMO)^50^ tool (v. 5.5.9) in the MEME suite was used to identify occurrences of ASCL1, POU2F1, and tandem POU2F1–ASCL1 motifs in the human genome (hg38). Significance thresholds of 1×10^-7^ (tandem POU2F1–ASCL1 motifs and “reversed” tandem POU2F1–ASCL1 motifs) and 1×10^-5^ (ASCL1 and POU2F1 motifs) were applied. Annotation of genomic loci containing each set of motifs was performed using HOMER^49^ v5.11 with default settings.

#### AlphaFold 3 prediction analysis

AlphaFold 3 (using Google’s AlphaFold 3 server)^52^ was used to generate 20 predictions of the structure of the ASCL1–POU2F1–TCF12 complex from 4 separate starting seeds. Predictions were generated using either a tandem POU2F1–ASCL1 motif (5’-**ATTTGCATA***CAGCTG*T-3’ and complement), a “reversed” tandem motif that swaps the relative positions of the POU2F1- and ASCL1-binding motifs (5’-*CAGCTG*T**ATTTGCATA**-3’ and complement), or without any DNA. Generated predictions were then examined in ChimeraX^68^ v1.11.1 and, using the “align” command, manually aligned to the structured domains of the highest-scoring model. For generation of the mean distance of residues/nucleotides from the mean aligned position, a Python script (written predominantly manually with some input by ChatGPT for the fixing of syntax errors and code annotation) was used to find the distance of each residue’s Cα (for proteins) or C1’ (for DNA) atom from the mean position of the atom across all twenty models, and repeated for all twenty models, to provide a measure of model consistency relative to the local environment. In addition, the corresponding pLDDT values for each Cα or C1’ atom from the AlphaFold 3 output JSON files were extracted and exported to a CSV file along with the distance data and manually input into GraphPad Prism for further analysis. ChimeraX was used to find potential hydrogen bonds and van der Waals contacts, with hydrogen bonds split into “strict” matches to ChimeraX’s hydrogen bonding criteria, and “relaxed” matches that allowed for a 1 Å increase to the distance cutoff and 35° to the angle cutoff, and van der Waals contacts were found using a VDW overlap of 0.4 Å in ChimeraX. A Python script was then used to extract the number of occurrences of “strict” hydrogen bonds, “relaxed” hydrogen bonds, and van der Waals contacts between each atom pair from the ChimeraX session, document the occurrence of each type of interaction between residue pairs across all twenty models (with only one allowed per residue, with “strict” hydrogen bonds taking priority, followed by “relaxed” hydrogen bonds, then van der Waals contacts), extract predicted aligned error (PAE) and estimated contact probability data from the AlphaFold 3 JSON outputs for each model, and output to a CSV file. Due to the large variability between predictions both within and across model seeds, especially with the predictions without DNA and the “reversed” motif predictions, all twenty outputs were used as standalone replicates. Welch’s unpaired t-test was used to find significant differences in model confidence and consistency of the DNA bases, as changes to the DNA sequences used in “reversed” motif predictions made pairwise comparisons unreliable.

#### Cloning and molecular biology

Oligonucleotides for primers and sgRNAs were ordered from SigmaAldrich. Pre-made vectors and gene cassettes were ordered from Addgene, and new DNA fragments were ordered as gBlocks from IDT (Integrated DNA Technologies). The Takara^®^ In-Fusion HD or Snap Assembly cloning kit was used to clone new plasmids according to manufacturer’s instructions. Plasmids were transformed into Stbl3^TM^ competent *E. coli* for antibiotic selection and plasmid amplification. All plasmid sequences were sequenced for validation prior to use.

#### Western blots and protein analysis

Cells were collected, washed 2x with room temperature PBS, counted for each sample and normalized by cell number. Equal numbers of cells were washed in PBS and resuspended in 1x Laemmli buffer, diluted in PBS, with 5% β-mercaptoethanol at a final concentration of ∼5×10^6^ cells/mL. Samples were boiled at 98°C for 15 minutes and stored at 4°C for a maximum of one month, or at −20°C for long-term storage. Immediately prior to loading on an SDS-PAGE gel, samples were boiled for 1 minute and centrifuged at 20,000 RCF for 5 minutes. Antibodies are included as Supplementary Table 8.

#### Statistics and Reproducibility

All statistical tests used to evaluate significance are detailed in the respective figure legends. All reported results were replicated across multiple experiments to generate reliable results, and replication is described in more detail in each figure legend. No statistical method was used to predetermine sample size. Sample size calculations were based on previously published data. Figure legends indicate the sample sizes for each set of experiments. All sample sizes were determined to be sufficient given that the differences among groups were consistent. No data was excluded from any experiment. Randomization was not performed for any experiments. Investigators were not blinded to allocation during experiments and outcome assessment.

## Supporting information

Uncropped Western Blots

## Data Availability

All genomic datasets are available at the GEO database under accession codes GSE317998 [reviewer token = anixowuiznohvot] (CRISPR screens), GSE317997 [reviewer token = urgbywaeppgnjgp] (RNA-seq), and GSE317996 [reviewer token = ktkxquwidbgffuj] (ChIP-seq). Raw uncropped Western Blots are published alongside this paper.

## Description of Supplementary Information

Supplementary Figure 1: *DLL3* and *POU2F1* are highly expressed in SCLC-A.

Supplementary Figure 2: ASCL1 activates *DLL3* and the neuroendocrine identity gene expression program in SCLC.

Supplementary Figure 3: POU2F1 co-occupies chromatin at some ASCL1 binding sites.

Supplementary Figure 4: Tandem POU2F1–ASCL1 motifs are enriched at cis-regulatory elements associated with neuroendocrine identity marker genes.

Supplementary Figure 5: AlphaFold 3 predictions decreased in confidence with a reversed tandem motif

Supplementary Figure 6: Tandem POU2F1–ASCL1 motifs inform POU2F1- and ASCL1-activated gene expression in small cell lung cancer cell lines.

Supplementary Table 1: Gene Expression in CCLE Cell Lines

Supplementary Table 2: CRISPR Screens Results

Supplementary Table 3: Gene Expression in Human small cell lung cancer (George et al. 2015)

Supplementary Table 4: *POU2F1* and *ASCL1* Knockout RNA-sequencing Results

Supplementary Table 5: Motif Analyses

Supplementary Table 6: Find Individual Motif Occurrences (FIMO) Results

Supplementary Table 7: Gene Sets for Gene Set Enrichment Analysis (GSEA)

Supplementary Table 8: Antibodies

## Acknowledgements

This study was supported by the Cold Spring Harbor Laboratory NCI Cancer Center Support under grant CA045508. C.R.V. was supported by the Pershing Square Sohn Cancer Research Alliance, and NCI grant CA290004. Additional funding was provided by Cold Spring Harbor Laboratory and Northwell Health Affiliation and Treeline Biosciences. P.J.C. was supported by NCI grant CA278591. D.S. was supported by NCI R50 grant CA305054. L.J. is an investigator of the Howard Hughes Medical Institute.

## Author Contributions

P.J.C. performed the formal analysis and wrote the initial draft of the manuscript. C.F. designed the methodology and carried out the investigation and validation studies, with assistance from D.S. in performing the marker-based CRISPR screens. J.B. performed the AlphaFold 3 analysis, and L.J. provided supervision and advice. O.K. and T.Y. subcloned the Brunello genome-wide sgRNA library. C.R.V. conceived the project, developed the methodology, supervised the research, acquired funding, and contributed to writing, editing, and review of the manuscript.

## Competing Interests

C.R.V. has received consulting fees from Flare Therapeutics, Roivant Sciences and C4 Therapeutics; has served on the advisory boards of KSQ Therapeutics, Syros Pharmaceuticals and Treeline Biosciences; has received research funding from Boehringer-Ingelheim and Treeline Biosciences; and owns stock in Treeline Biosciences. The remaining authors declare no competing interests.

