## Supplementary figures and images for "Marker-based CRISPR screens identify POU2F1 as a regulator of DLL3 and neuroendocrine identity in small cell lung cancer"

### Uncropped Western Blots

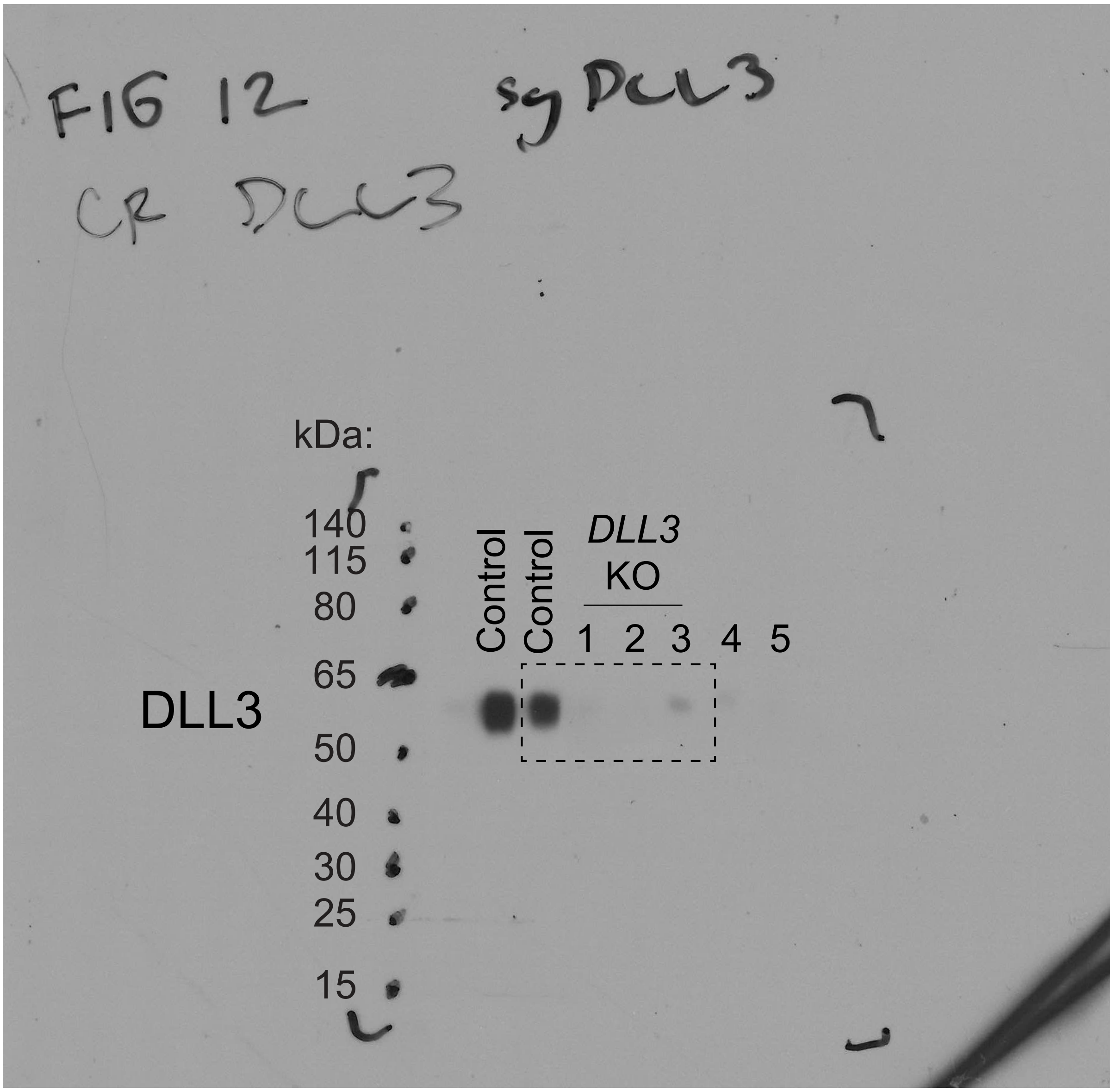

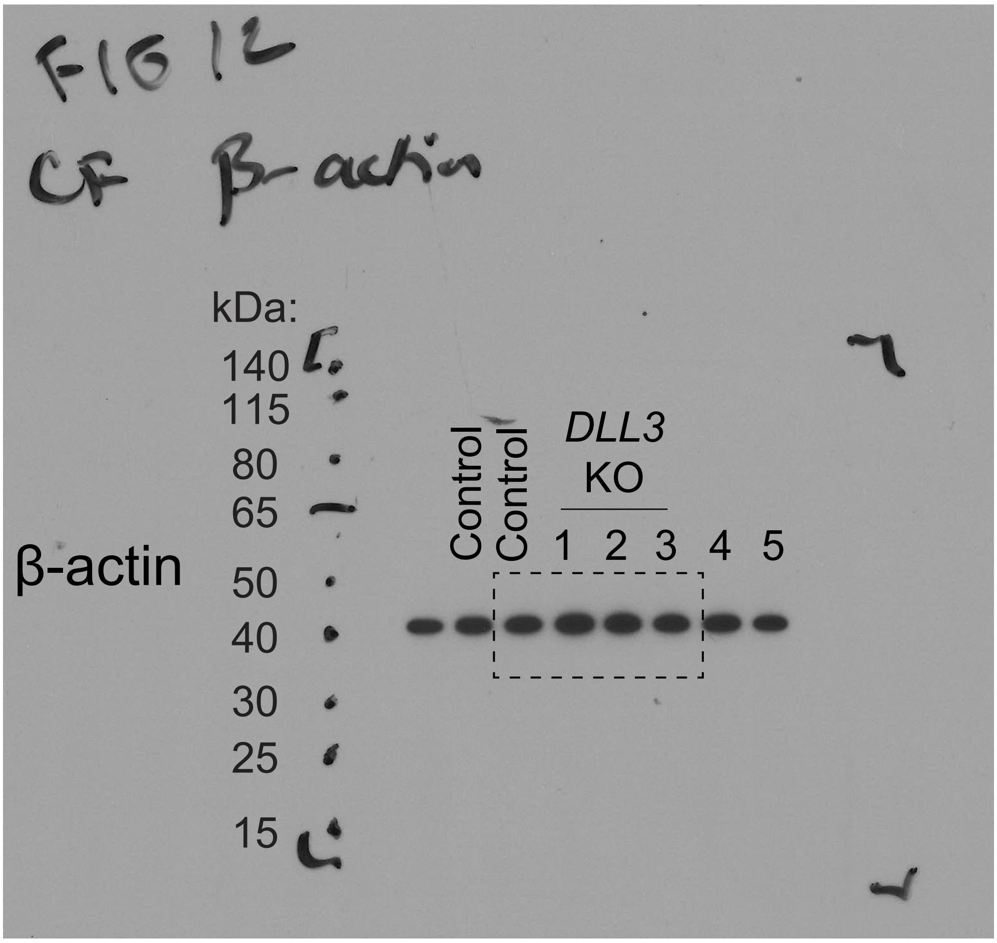

Figure 2A NCI-H209

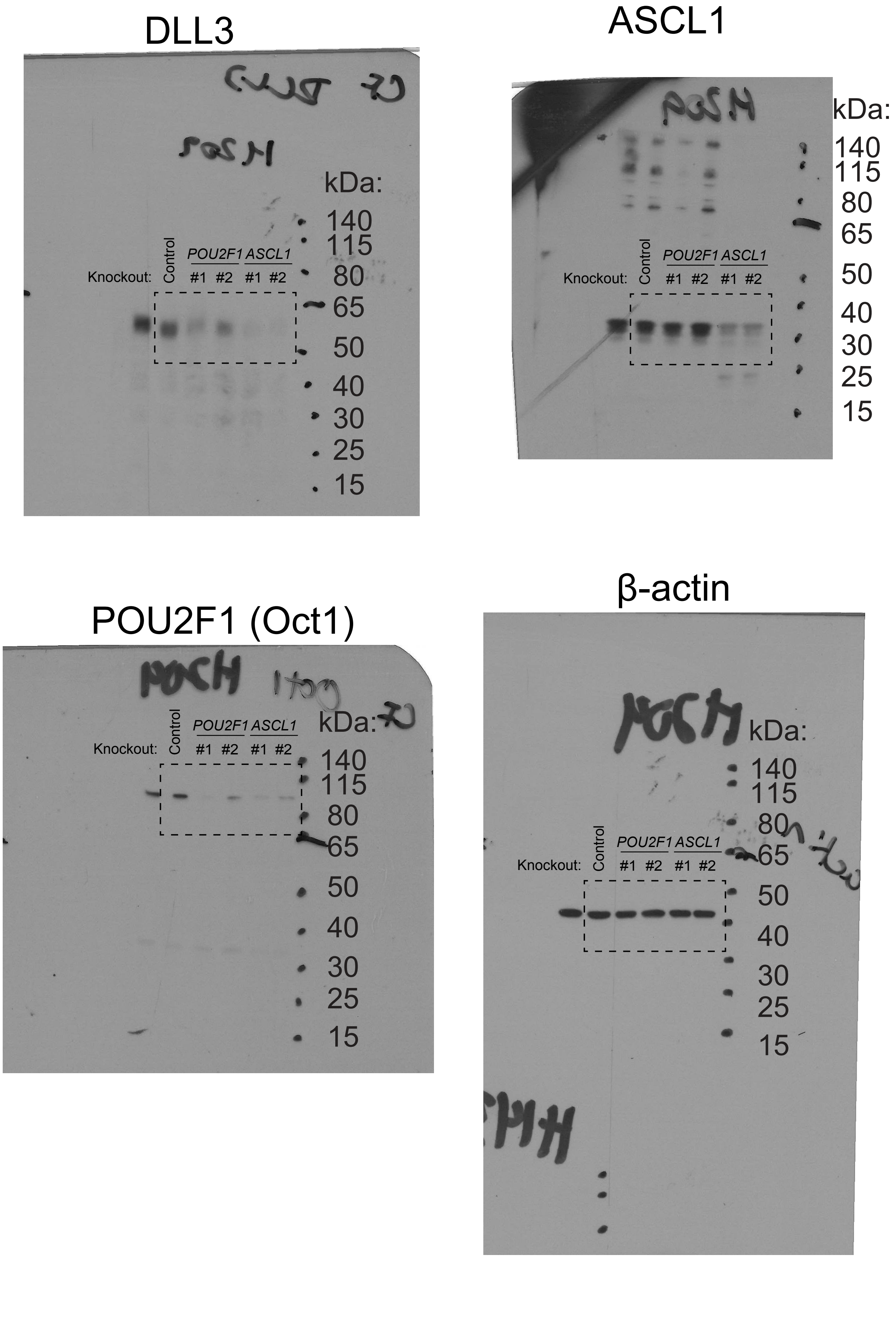

#### Figure 2A

NCI-H1436

DLL3

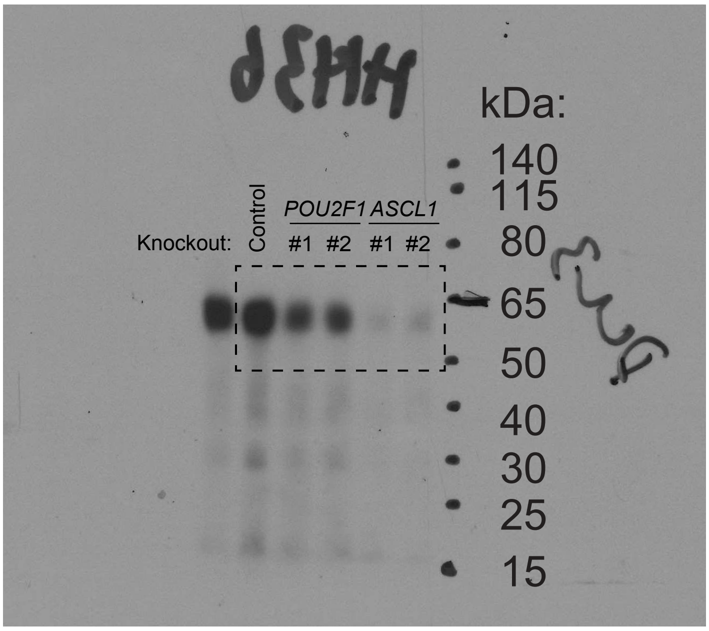


ASCL1

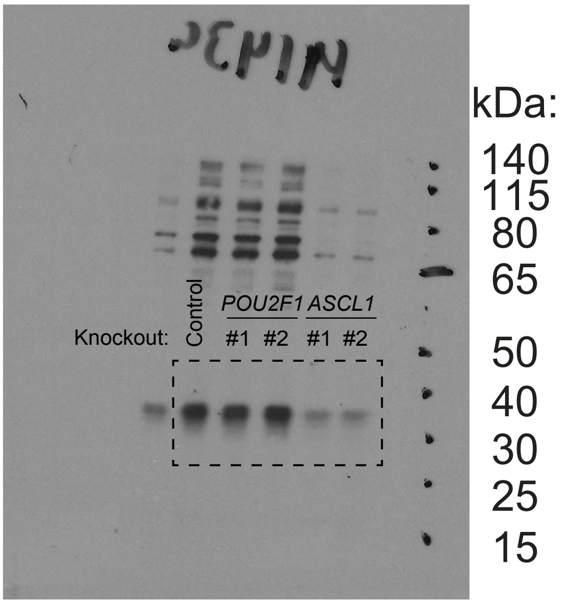


POU2F1 (Oct1)

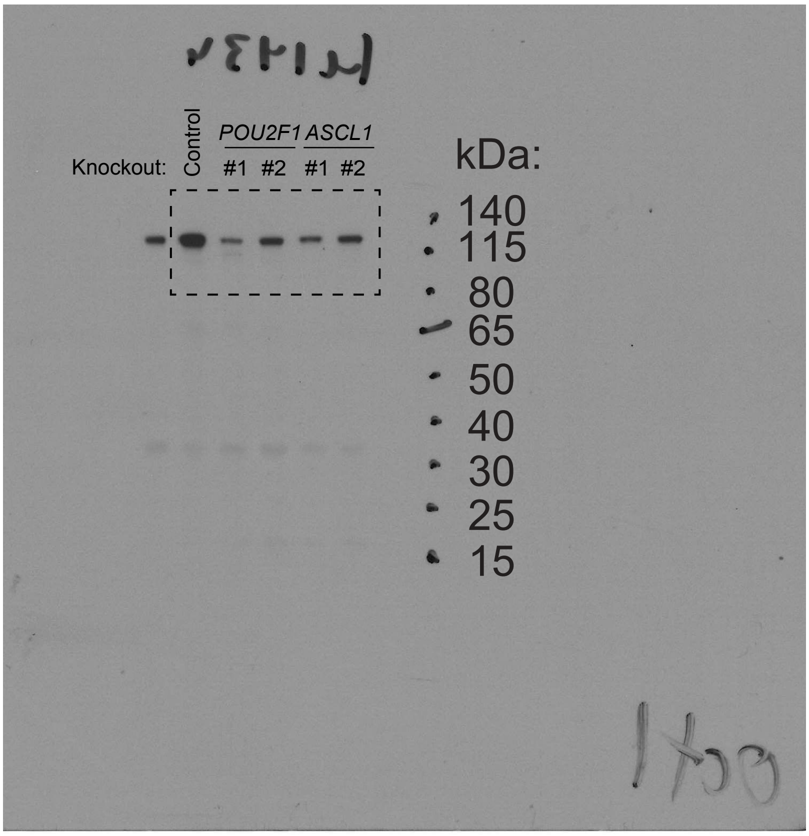


β-actin

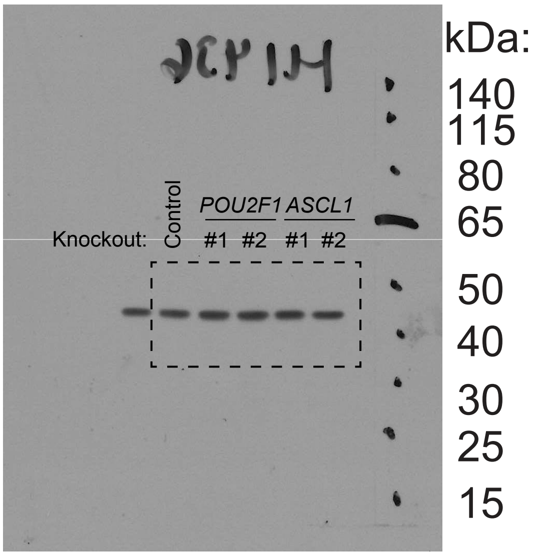

Figure 2A NCI-H1836

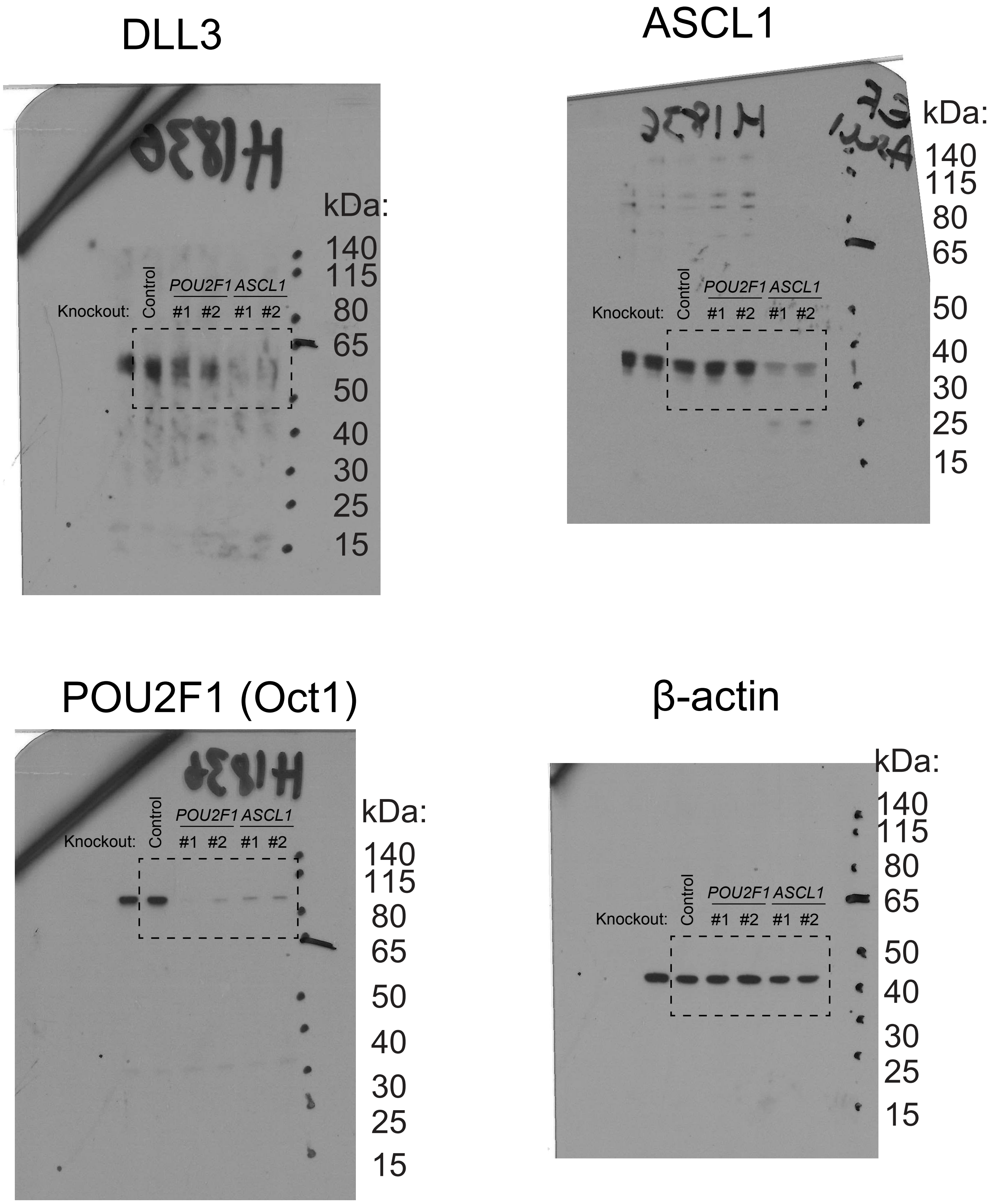
