## Supplementary Figures 1-6 for "Marker-based CRISPR screens identify POU2F1 as a regulator of DLL3 and neuroendocrine identity in small cell lung cancer"

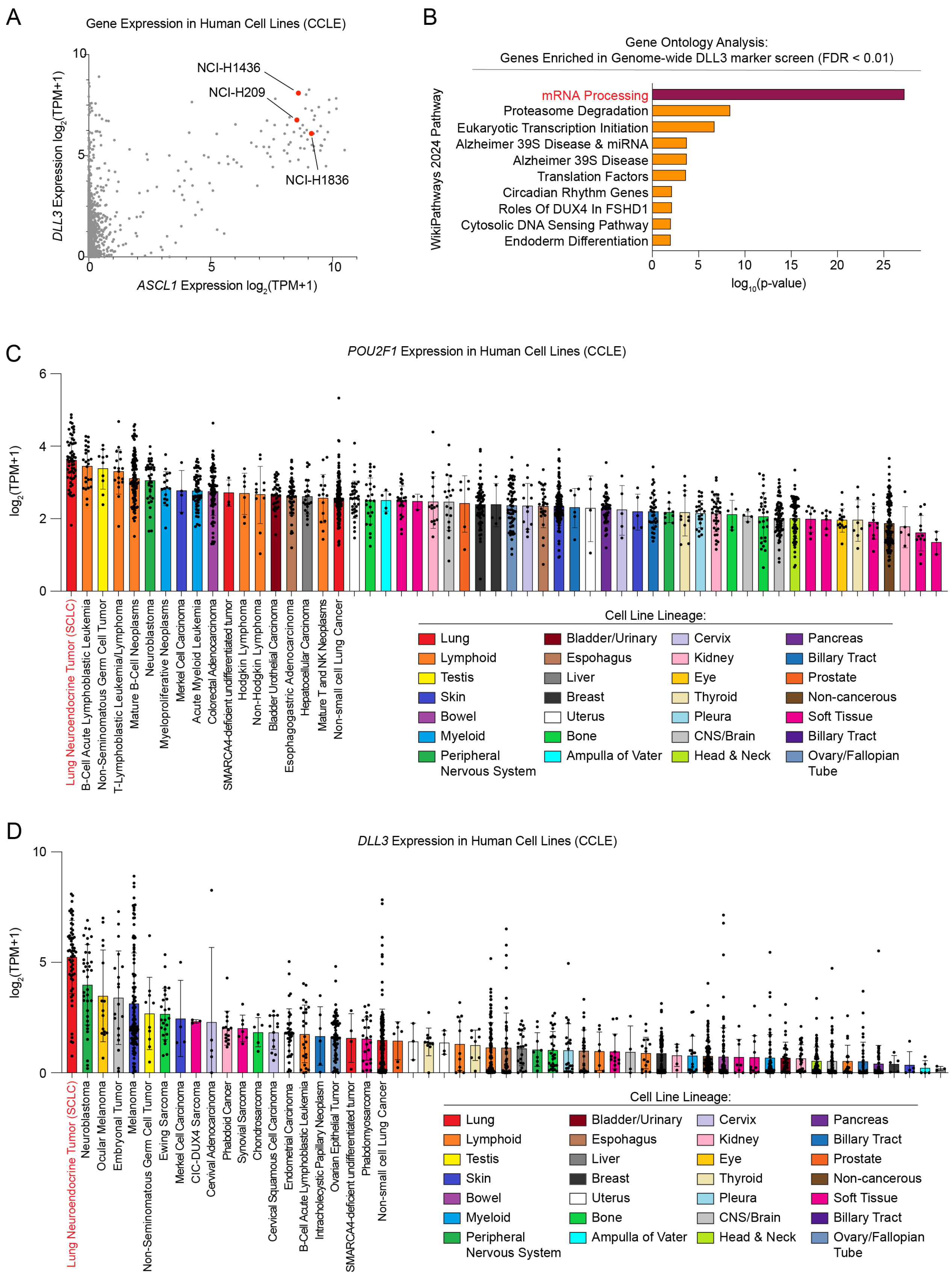


**Supplementary Figure 1: *DLL3* and *POU2F1* are highly expressed in SCLC-A**. (A,C,D) Gene expression in human cell lines (CCLE). TPM = transcripts per million (A) Expression of *ASCL1* and *DLL3* in human small cell lung cancer cell lines. The NCI-H209, NCI-H1436, and NCI-H1836

cell lines are highlighted and labeled. (B) Enrichr gene ontology analysis of 129 genes enriched in the DLL3$^{low}$ population of the genome-wide DLL3 marker-based CRISPR screen (FDR $< 0.01$). The top 10 WikiPathway signatures are ranked by their significance (p-value) and the most significant terms ($p < 0.05$) are highlighted. (C-D) POU2F1 (C) and DLL3 (D) expression is plotted as log transformed TPM values. Each point = 1 cell line. Bars represent the mean ± SD. Bars are colored according to cell line lineage, and rank ordered by mean expression. Lineages with > 1 cell line are included.

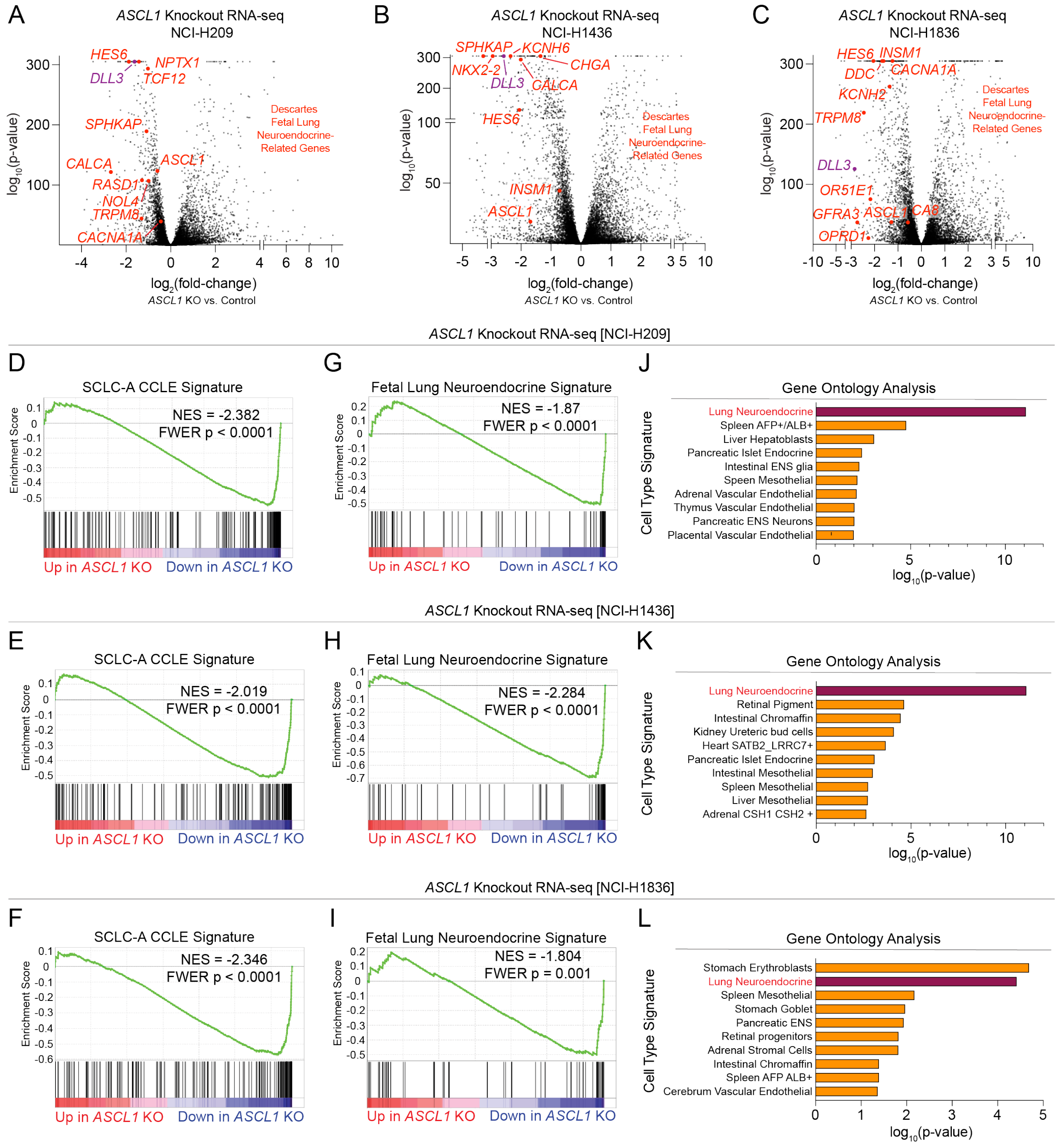


**Supplementary Figure 2: ASCL1 activates *DLL3* and the neuroendocrine identity gene expression program in SCLC**. (A-L) RNA sequencing in NCI-H209 (A,D,G,J), NCI-H1436 (B,E,H,K), and NCI-H1836 (C,F,I,L) on day 4 (A,D,G,J) or day 6 (B,C,E,F,H,I,K,L) following CRISPR-Cas9 knockout (KO) of *ASCL1*, or *ROSA26* (control). Fold change and significance calculated by DESeq2. n = 6 biological replicates per timepoint, per cell line. (A-C) Volcano plots of differentially expressed genes following *ASCL1* KO. *DLL3* and additional genes related to fetal lung neuroendocrine identity (DESCARTES database; Cao et al. 2021) are labeled. (D-I) Gene set enrichment analysis (GSEA) of differentially expressed genes following *ASCL1* KO. Significance calculated by GSEA. NES = Normalized Enrichment score. FWER = Family-wise Error Rate. (J-L) Enrichr Gene ontology analysis of top 250 downregulated genes following *ASCL1* KO. Top 10 DESCARTES Cell Type signatures are ranked ordered by their significance (p-value) and the most significant terms ($p < 0.05$) are highlighted.

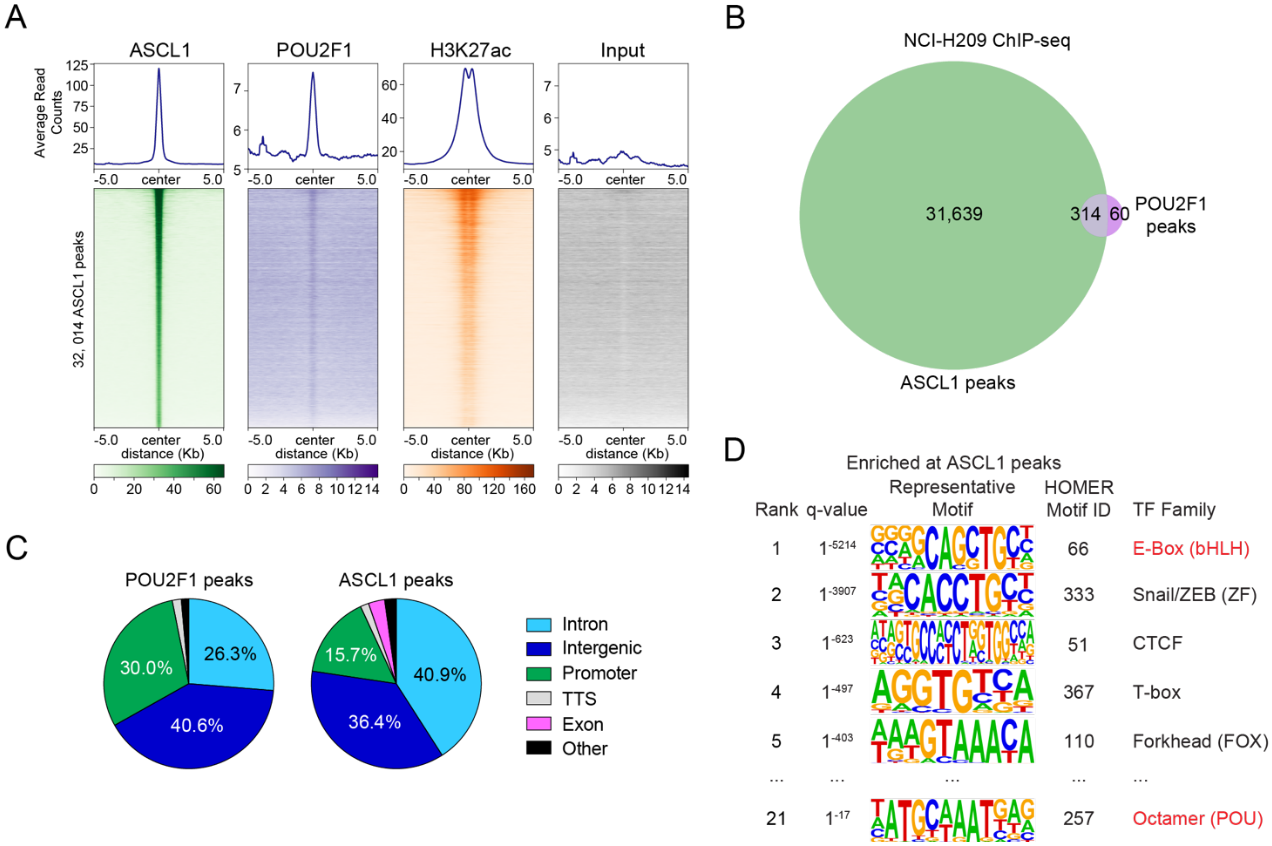


**Supplementary Figure 3: POU2F1 co-occupies chromatin at some ASCL1 binding sites**. (A-D) ASCL1, POU2F1, and H3K27ac ChIP-seq in NCI-H209 (A) Heatmaps for ASCL1, POU2F1, H3K27ac ChIP-seq and input (control) at all ASCL1 peaks. Rows = 10kb genomic regions centered on an ASCL1 peak summit. Rows are ordered by ASCL1 signal, and this ordering is applied to all heatmaps. Metagene plots show the average signal for each factor across all peaks, plotted above each heatmap. (B) Venn diagram of called ASCL1 and POU2F1 peaks (MACS2 $q < 0.01$). (C) HOMER annotations for 31,974 ASCL1 peaks and 374 consensus POU2F1 peaks. (D) HOMER motif enrichment analysis of 31,953 called ASCL1 peaks (MACS2 $q < 0.01$). The top 5 transcription factor family motifs were selected.

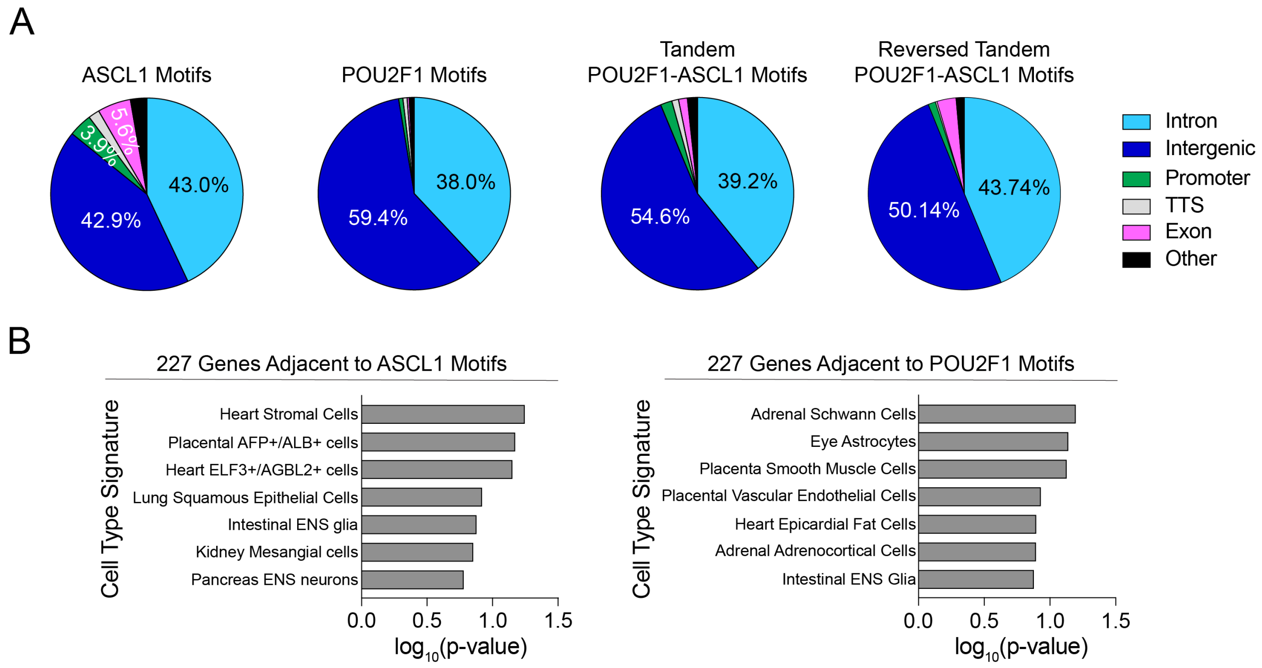


**Supplementary Figure 4: Tandem POU2F1–ASCL1 motifs are enriched at cis-regulatory elements associated with neuroendocrine identity marker genes.** (A) HOMER annotations for 21,824 loci containing an ASCL1 motif (FIMO; $p < 1x10^5$), 90.253 loci containing a POU2F1 peak (FIMO; $p < 1x10^5$), 876 loci containing a Tandem POU2F1–ASCL1 motif (FIMO; $p < 1x10^7$), and 718 loci containing a reversed Tandem POU2F1–ASCL1 motif (FIMO; $p < 1x10^7$) (hg38). (B) Enrichr Gene ontology analysis of 227 randomly selected, annotated genes within 20kb of an ASCL1 ($p < 1x10^5$), or POU2F1 ($p < 1x10^5$) motif (hg38). The top 7 DESCARTES Cell Type signatures are ranked ordered by their significance (p-value).

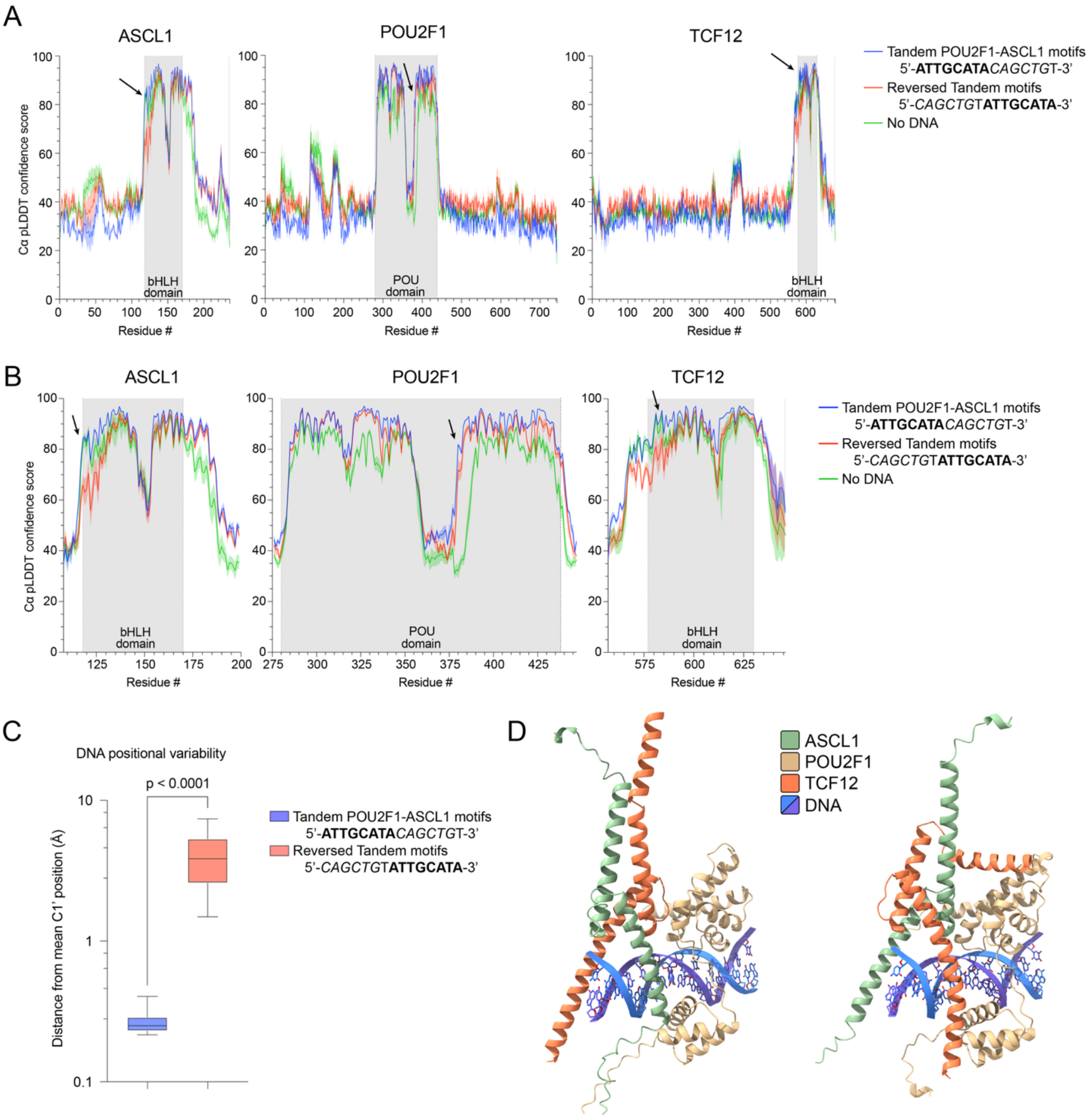


**Supplementary Figure 5: AlphaFold 3 predictions decreased in confidence with a reversed tandem motif.** (A) A plot of the mean Cα pLDDT confidence scores across all residues of ASCL1, POU2F1, or TCF12, with the standard deviation surrounding it. **Bold letters** were used to represent the POU2F1 binding motif, while *italics* indicate the ASCL1 binding motif. Arrows point to regions of predicted protein-protein interactions: the N-terminal region adjacent to the bHLH of ASCL1, residues 377–382 of POU2F1, and the N-terminus of the TCF12 bHLH, respectively. (B) Similar to (A), but focused on windows surrounding the structured domains of each protein. (C) Distances of the C1' atoms of each DNA nucleotide (N=32 each) from the mean C1' position in AlphaFold 3 models containing the ASCL1–POU2F1 tandem motif or a "reversed" motif. Smaller distances from the mean aligned position indicate greater positional consistency across models. Each data point was generated from the mean distance of the C1' atom from its mean aligned position across all models. Boxes represent the mean ± interquartile range (IQR). Error bars extend to the minimum and maximum values. Statistical significance was evaluated using Welch's unpaired t-test. (D) Two distinct conformations are seen in the "reversed" tandem motif ASCL1–

POU2F1–TCF12 AlphaFold 3 predictions; the predominant conformation seen in 15/20 models (right), and one with an inversion of the ASCL1–TCF12 heterodimer orientation relative to POU2F1 and the DNA (left) seen in 5/20 models, indicating heterogeneity of predictions at a protein domain level.

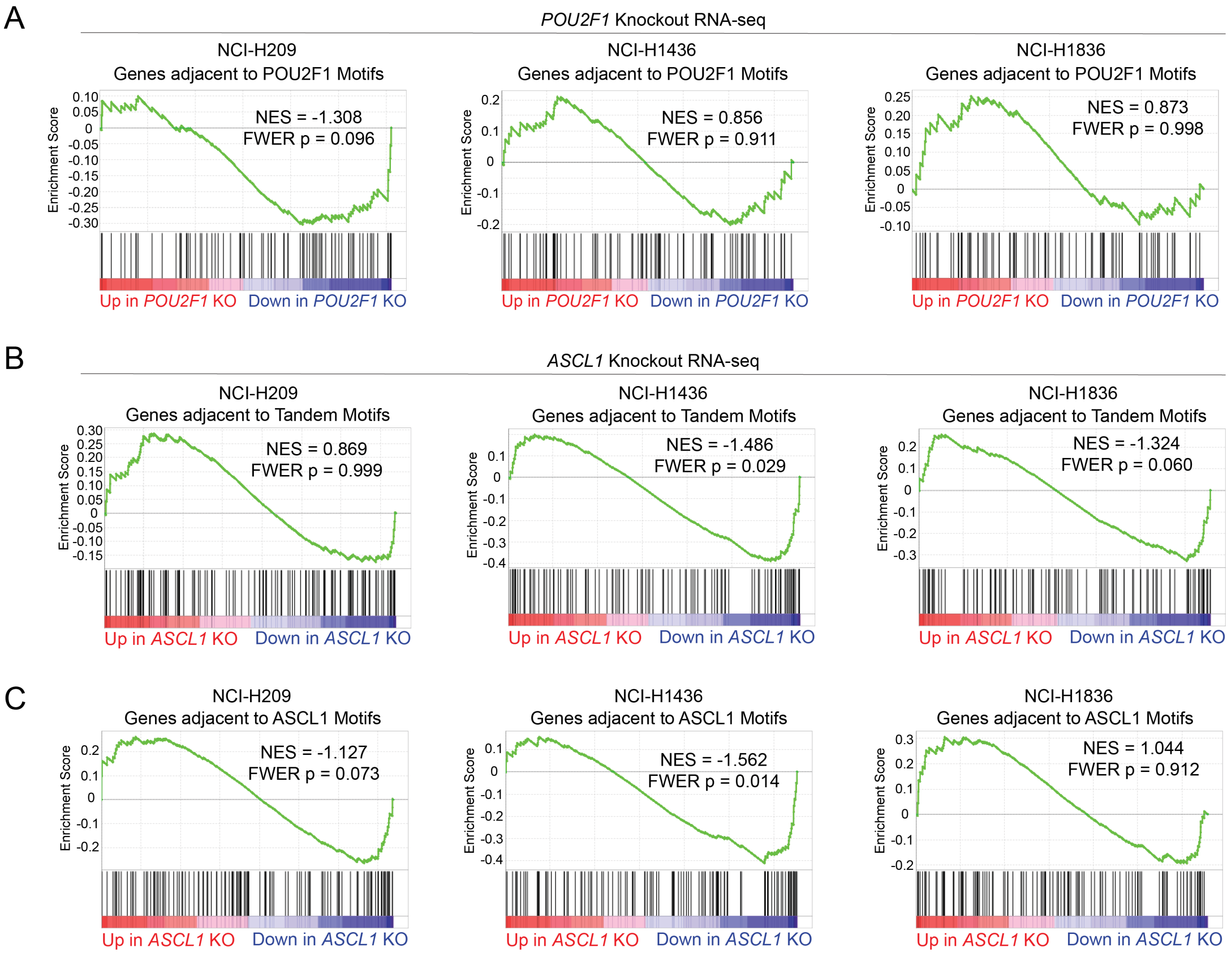


**Supplementary Figure 6: Tandem POU2F1–ASCL1 motifs inform POU2F1- and ASCL1-activated gene expression in small cell lung cancer cell lines.** (A-C) RNA sequencing in NCI-H209, NCI-H1436, and NCI-H1836 following CRISPR-Cas9 knockout (KO) of *POU2F1*, *ASCL1*, or *ROSA26* (control). Fold change and significance calculated by DESeq2. n = 6 biological replicates per timepoint, per cell line. Gene set enrichment analysis (GSEA) of differentially expressed genes following *POU2F1* or *ASCL1* KO. NES = Normalized Enrichment score. FWER = Family-wise Error Rate.
